# The contribution of sex chromosome conflict to disrupted spermatogenesis in hybrid house mice

**DOI:** 10.1101/2022.07.19.499960

**Authors:** Emily E. K. Kopania, Eleanor M. Watson, Claudia C. Rathje, Benjamin M. Skinner, Peter J. I. Ellis, Erica L. Larson, Jeffrey M. Good

## Abstract

Incompatibilities on the sex chromosomes are important in the evolution of hybrid male sterility, but the evolutionary forces underlying this phenomenon are unclear. House mice (*Mus musculus*) lineages have provided powerful models for understanding the genetic basis of hybrid male sterility. X chromosome-autosome interactions cause strong incompatibilities in *Mus musculus* F1 hybrids, but variation in sterility phenotypes suggests a more complex genetic basis. Additionally, XY chromosome conflict has resulted in rapid expansions of ampliconic genes with dosage-dependent expression that is essential to spermatogenesis. Here we evaluated the contribution of XY lineage mismatch to male fertility and stage-specific gene expression in hybrid mice. We performed backcrosses between two house mouse subspecies to generate reciprocal Y-introgression strains and used these strains to test the effects of XY mismatch in hybrids. Our transcriptome analyses of sorted spermatid cells revealed widespread overexpression of the X chromosome in sterile F1 hybrids independent of Y chromosome subspecies origin. Thus, postmeiotic overexpression of the X chromosome in sterile F1 mouse hybrids is likely a downstream consequence of disrupted meiotic X-inactivation rather than XY gene copy number imbalance. Y-chromosome introgression did result in subfertility phenotypes and disrupted expression of several autosomal genes in mice with an otherwise nonhybrid genomic background, suggesting that Y-linked incompatibilities contribute to reproductive barriers, but likely not as a direct consequence of XY conflict. Collectively, these findings suggest that rapid sex chromosome gene family evolution driven by genomic conflict has not resulted in strong male reproductive barriers between these subspecies of house mice.

## Introduction

Sex chromosomes are often involved in the evolution of reproductive isolation between animal species (Coyne and Orr 1989; Turelli and Orr 2000; Presgraves and Meiklejohn 2021), with hybrid sterility or inviability arising more often in the heterogametic sex (i.e., Haldane’s Rule, Haldane 1922; Coyne and Orr 2004). Hybrid incompatibilities also tend to accumulate more rapidly on the X chromosome (Masly and Presgraves 2007), which is referred to as the large X-effect (Coyne and Orr 1989). Known as the two rules of speciation (Coyne and Orr 1989; Coyne and Orr 2004), these patterns have been supported across diverse taxa (Good et al. 2008a; Davis et al. 2015; Bi et al. 2019; Matute and Cooper 2021; Presgraves and Meiklejohn 2021) and undoubtedly drive the early stages of intrinsic reproductive isolation in many systems. Both Haldane’s rule and the large X-effect appear particularly strong when considering hybrid male sterility in XY systems, suggesting an important role for X chromosome evolution in both speciation and the evolution of spermatogenesis. However, it remains unclear to what extent these general patterns reflect common evolutionary processes, functional mechanisms unique to sex chromosomes, or a mixture of both (Meiklejohn and Tao 2010).

Intrinsic reproductive barriers between nascent species often arise as an indirect consequence of rapid evolution within populations (Dobzhansky 1937; Coyne and Orr 2004; Coughlan and Matute 2020), so the outsized contribution of sex chromosomes to male sterility may be an inevitable consequence of rapid molecular evolution on the X and Y chromosomes. For example, recurrent genomic conflict is thought to be rampant on the X and Y chromosomes because selfish genetic elements are more likely to arise on sex chromosomes (i.e., meiotic drive *sensu lato*; Frank 1991; Hurst and Pomiankowski 1991; Meiklejohn and Tao 2010; Lindholm et al. 2016). Hemizygosity of the X chromosome is also expected to promote more rapid adaptive molecular evolution relative to the autosomes across a broad range of conditions (i.e., the faster-X effect; Charlesworth et al. 1987; Vicoso and Charlesworth 2009). Note that hemizygosity on the X and Y chromosomes will also result in differential exposure of hybrid incompatibilities on the sex chromosomes in males if incompatibilities tend to be at least partially recessive (Turelli and Orr 1995; Turelli and Orr 2000). However, progress on understanding how often these diverse evolutionary processes contribute to the evolution of hybrid male sterility has been hampered by a lack of data on the genetic underpinnings of reproductive isolation.

From a mechanistic perspective, the X and Y chromosomes are also subject to unique regulatory processes during mammalian spermatogenesis that are critical for normal male fertility and shape patterns of molecular evolution (Larson, et al. 2018a). Both the X and Y chromosomes are packaged into condensed chromatin early in meiosis, resulting in transcriptional silencing of most sex-linked genes known as meiotic sex chromosome inactivation (MSCI; McKee and Handel 1993). Repressive chromatin persists through the postmeiotic stages (Namekawa, et al. 2006), although many essential X- and Y-linked genes are highly expressed in postmeiotic, haploid round spermatids (Mueller, et al. 2008; Sin and Namekawa 2013). Failure to broadly repress X-linked expression during these critical meiotic and postmeiotic stages can trigger spermatogenic disruption, reduced sperm production, and abnormal sperm morphology (Burgoyne et al. 2009; Turner 2015). Interestingly, sex chromosome repression during both stages appears prone to disruption in hybrid mammals (Mihola et al. 2009; Good et al. 2010; Campbell et al. 2013; Davis et al. 2015; Larson et al. 2017), which may reflect common regulatory pathways underlying the evolution of hybrid male sterility (Bhattacharyya et al. 2013; Larson et al. 2021). Understanding how these intermediate developmental sterility phenotypes relate to genomic conflict and the broader evolutionary dynamics of the sex chromosomes awaits more data.

House mice (*Mus musculus*) have emerged as predominant models for understanding both the basic molecular control of spermatogenesis and the evolution of hybrid male sterility in mammals (Phifer-Rixey and Nachman 2015). Closely related subspecies of mice, *Mus musculus musculus* and *M. m. domesticus* (hereafter, “*musculus*” and “*domesticus*”), readily hybridize in both the lab and along a natural hybrid zone in Europe (Janoušek et al. 2012). Hybrid male sterility is the strongest and likely primary reproductive barrier isolating these incipient species in nature (Vyskočilová, et al. 2005; Turner, et al. 2012) and in the lab (Good et al. 2008b; Vyskočilová et al. 2009) following Haldane’s rule (Haldane 1922; but see Suzuki and Nachman 2015). Male sterility is polymorphic with laboratory crosses yielding sterile, subfertile, or fertile male hybrids depending on genotype and cross direction (Good et al. 2008b; Balcova et al. 2016; Larson et al. 2018b; Widmayer et al. 2020); *musculus*^♀^ × *domesticus*^♂^ crosses usually result in sterile F1 males, while the reciprocal cross tends to be more fertile (Good et al. 2008b). This asymmetry is caused by epistatic incompatibilities that are exposed on the *musculus* X chromosome in hybrid males (Storchová et al. 2004; Good et al. 2008a; Turner and Harr 2014). House mice also remain the only mammalian system where the evolution of a specific gene, *Prdm9*, has been directly linked to the evolution of intrinsic reproductive barriers (Mihola et al. 2009; Bhattacharyya et al. 2013; Mukaj et al. 2020). *Prdm9* is an autosomal gene encoding a DNA-binding protein that directs double stranded breaks where meiotic recombination occurs (Grey et al. 2011). PRDM9 binding sites evolve rapidly (Oliver et al. 2009; Baker et al. 2015), leading to asymmetric binding in hybrid mice that triggers autosomal asynapsis and disruption of MSCI during early pachytene of Meiosis I (Mihola et al. 2009; Davies et al. 2016). *Prdm9*-related sterility depends on *Prdm9* heterozygosity and epistatic interactions with other unlinked factors, including a major incompatibility locus, *Hstx2*, located near the middle the *musculus* X chromosome (Forejt et al. 2021). This same X-linked region also influences hybrid male sterility in backcrossed consomic models (i.e., presumably independent of *Prdm9*; Storchová et al. 2004; Good et al. 2008a), and recombination rate variation between *M. m. musculus* and another subspecies, *M. m. castaneus* (Dumont and Payseur 2011).

This broad foundation on the genetics of hybrid male sterility provides an opportunity to further unravel the various evolutionary and mechanistic processes that contribute to the large X-effect in mice. *Prdm9*-related sterility plays a central role in the evolution of hybrid male sterility and the disruption of MSCI in F1 mouse hybrids (Forejt et al. 2021; Larson et al. 2021). However, X- and Y-linked hybrid sterility arises across a broader range of genetic architectures and phenotypes than can be easily ascribed to *Prdm9*-related interactions (Campbell et al. 2012; Campbell and Nachman 2014; Larson et al. 2018b; Larson et al. 2021). The mouse X and Y chromosomes also contain clusters of several high copy ampliconic genes (Mueller et al. 2008; Soh et al. 2014; Case et al. 2015; Morgan and Pardo-Manuel De Villena 2017; Larson et al. 2021) that appear to have evolved in response to intense intragenomic conflict (Cocquet et al. 2009; Ellis et al. 2011; Cocquet et al. 2012). These X- and Y-linked gene clusters are primarily expressed in postmeiotic cells with repressed sex chromatin (Namekawa et al. 2006; Sin et al. 2012) and thus increases in copy number may help counteract repressive chromatin (Ellis et al. 2011; Mueller et al. 2013; Sin and Namekawa 2013). Conflict arises because the maintenance of repressive postmeiotic sex chromatin appears to be controlled by dosage dependent interactions between X-linked (*Slx* and *Slxl1*) and Y-linked (*Sly*) gene families (Cocquet et al. 2012; Kruger et al. 2019). Experimental knockdowns of *Slx* and *Slxl1* showed increased sex chromosome repression, abnormal sperm head morphology, and an excess of male offspring. In contrast, knockdowns of *Sly* showed sex chromosome overexpression, abnormal sperm head morphology, and an excess of female offspring (Cocquet et al. 2009; Cocquet et al. 2012) due to reduced motility of Y-bearing sperm (Rathje et al. 2019). CRISPR-based deletions have further shown that sex-ratio distortion is primarily mediated by *Slxl1* versus *Sly* competition for the spindlin proteins (SPIN1, SSTY1/2; Kruger et al. 2019).

Copy numbers of *Slx, Slxl1*, and *Sly* genes have co-evolved in different mouse lineages (Ellis et al. 2011; Good 2012; Morgan and Pardo-Manuel De Villena 2017), such that hybrids could have copy number mismatch sufficient to generate dosage-based sterility phenotypes seen in genetic manipulation studies (Ellis et al. 2011). In support of this model, hybrid interactions between the *musculus* X and the *domesticus* Y have been shown to cause abnormal sperm head morphology (Campbell et al. 2012; Campbell and Nachman 2014), and male sterility is associated with extensive overexpression of the sex chromosomes in postmeiotic round spermatids in *musculus*^♀^ × *domesticus*^♂^ mice (Larson et al. 2017). These hybrids have proportionally higher numbers of *Slx* and *Slxl1* relative to *Sly* copies compared to nonhybrids and show patterns qualitatively consistent with the overexpression phenotypes observed in *Sly* knockdown and *Slx*/*Slxl1* duplication mice (Cocquet et al. 2012; Kruger et al. 2019). However, postmeiotic sex chromatin repression is thought to partially depend on repressive histone marks established during meiosis (Turner et al. 2006), and the same direction of the hybrid cross also shows disrupted MSCI in meiotic spermatocytes (Campbell et al. 2013; Larson et al. 2017). Thus, it remains unclear if the disruption of repressive postmeiotic chromatin is a consequence of XY mismatch or primarily a downstream epigenetic effect of deleterious interactions between the *musculus* X chromosome and *Prdm9* during meiosis (Larson et al. 2021).

Here, we advance understanding of the basis of hybrid male sterility in this system using a reciprocal backcrossing scheme to generate mice with the Y chromosome of one *Mus musculus* subspecies on the genomic background of another (Figure 1A). We used these Y-consomic genetic models to perform two reciprocal cross experiments while controlling for the effects of inbreeding. First, we tested for the potential rescue of sterility phenotypes in hybrid males with F1 autosomal genotypes but with matching X and Y chromosomes from the same subspecies (Hybrid F1 XY Match; Figure 1B). This experiment allowed us to tease apart XY interactions (i.e., *Slx* and *Slxl1* versus *Sly*) from X-autosomal interactions (i.e., *Prdm9*-related sterility). Second, we tested the effects of XY mismatch on different subspecific backgrounds (Nonhybrid XY Mismatch; Figure 1B). This experiment allowed us to test for incompatibilities exposed on introgressed Y chromosomes that occur independently of other hybrid interactions. We used genome sequencing to quantify X- and Y-linked gene copy numbers, quantified male reproductive phenotypes (testis weight and high-resolution sperm head morphology), and used Fluorescence-Activated Cell Sorting (FACS) to isolate cell populations enriched for either early meiotic leptotene-zygotene spermatocytes or postmeiotic round spermatids. We used these experiments to address three main questions: (i) Does XY mismatch cause abnormal male reproductive traits? (ii) Do differences in copy number predict differences in ampliconic gene family expression levels during late spermatogenesis? (iii) Is XY mismatch associated with disrupted gene expression during late spermatogenesis, particularly on the sex chromosomes?

**Figure 1.**
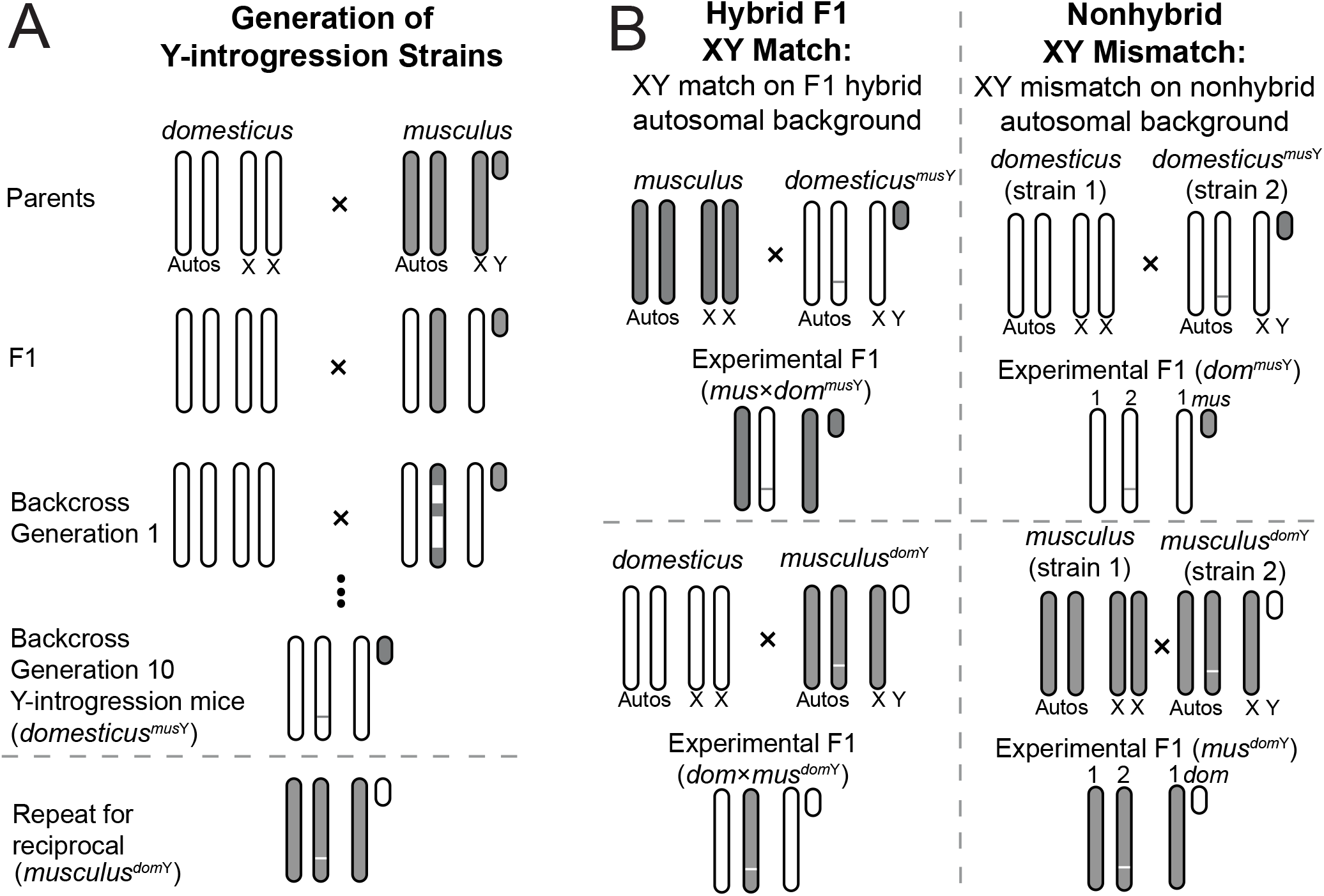
Experimental design. (A) Backcrosses used to generate Y-introgression mouse strains. We performed 10 generations of backcrosses in reciprocal directions to generate mice with a *Mus musculus domesticus* (*domesticus*) genetic background and *Mus musculus musculus* (*musculus*) Y chromosome (*domesticus*^*mus*Y^) and mice with a *musculus* genetic background and *domesticus* Y chromosome (*musculus*^*dom*Y^). The thin horizontal line on the autosomes represents residual autosomal introgression, which is theoretically expected to represent about 0.1% of the autosomes. (B) Crosses were performed with Y-introgression mice to produce two types of experimental F1 mice. For Hybrid F1 XY Match, we crossed Y-introgression males to females from the other subspecies to generate F1 mice with hybrid autosomes but matched sex chromosomes. For Nonhybrid XY Mismatch, we crossed Y-introgression males to females from a different strain but the same subspecies to generate F1 mice with XY mismatch and nonhybrid autosomes. Autos = autosomes, X = X chromosome, Y = Y chromosome.

## Materials and Methods

### Mouse resources and experimental design

We sought to test the effects of XY mismatch independent of the effects of X-autosomal incompatibilities and inbreeding. To do so, we conducted two experiments: (1) a “Hybrid F1 XY Match” experiment to test if matching the subspecies origin of the X and Y rescued expression and reproductive phenotypes on an otherwise F1 hybrid autosomal background, and (2) a “Nonhybrid XY Mismatch” experiment to test if introgressed XY subspecies origin mismatch disrupted expression and reproductive phenotypes on a nonhybrid autosomal background. To breed mice for these experiments, we first generated reciprocal consomic introgression strains with the Y chromosome from one subspecies on the genetic background of the other by backcrossing *musculus* (PWK) and *domesticus* (LEWES) for 10 generations, which we refer to as *musculus*^*dom*Y^ and *domesticusmus*Y (Figure 1A). We tested to ensure our Y-introgression strains had copy number mismatch representative of that expected in natural hybrids. We used publicly available whole genome sequence data to estimate copy number in wild house mice (PRJEB9450 for *domesticus*, 14 males and 1 female, Pezer et al. 2015; PRJEB11742 for *musculus*, 5 males and 11 females, Harr et al. 2016) and wild-derived inbred laboratory mouse strains representing *musculus* (PWK/ PhJ and CZECHII/EiJ) and *domesticus* (LEWES/EiJ and WSB/EiJ; PRJNA732719; one male individual per strain; Larson et al. 2021). We then used these Y-introgression strains to perform two experiments and test the effects of XY mismatch on hybrid sterility independent of X-autosomal incompatibilities (Figure 1B).

#### Experiment 1, Hybrid F1 XY Match

To test the effects of X-autosomal F1 incompatibilities without the effect of sex chromosome mismatch, we crossed Y-introgression males to females with the same autosomal and X chromosome type as the male Y chromosome (LEWES or PWK). This generated mice with an F1 hybrid autosomal background and X-autosomal mismatch but X and Y chromosomes from the same subspecies. Throughout the text, we refer to these mice as *mus*×*dommus*Y and *dom*×*mus*^*dom*Y^. We compared these mice to standard F1 hybrid mice with the same X chromosome and autosomal background but no Y chromosome introgression (PWK^♀^ × LEWES^♂^, hereafter “*mus*×*dom*” and LEWES^♀^ × PWK^♂^, hereafter “*dom*×*mus*”).

#### Experiment 2, Nonhybrid XY Mismatch

To test the effects of XY mismatch while controlling for inbreeding effects, we crossed Y-introgression males to females from the same subspecies but a different strain from the genomic background of the Y-introgression strain (CZECHII or WSB). This generated mice with a nonhybrid (intrasubspecific) F1 autosomal background and mismatched sex chromosomes (i.e., no X-autosomal mismatch), which we will refer to as *mus*^*dom*Y^ and *dom*^*mus*Y^. We compared these to intrasubspecific F1 mice with the same autosomal background as these F1 Y-introgression mice, but without sex chromosome mismatch (CZECHII^♀^ × PWK^♂^, hereafter “*mus*” and WSB^♀^ × LEWES^♂^, hereafter “*dom*”). Note that these Nonhybrid XY Mismatch mice had X chromosomes from different laboratory strains than the Hybrid F1 XY Match mice of the same subspecies as a necessary consequence of breeding mice with a heterozygous F1 background.

All mice from wild-derived inbred strains, Y-introgression strains, and experimental crosses were maintained in breeding colonies at the University of Montana (UM) Department of Laboratory Animal Resources (IACUC protocols 002-13, 050-15, and 062-18), which were initially purchased from The Jackson Laboratory, Bar Harbor, ME in 2010. Replacement stock of LEWES/EiJ mice were ordered in 2013, and these mice were used for the backcrosses to generate the *dom*^*mus*Y^ Y-introgression strains, as dames in the *dom* intrasubspecific F1s, and as sires in the *dom*×*mus* and *dom*×*mus*^*dom*Y^ crosses.

### Whole genome sequencing and copy number estimation

We sequenced whole genomes from one male mouse of each Y-introgression strain to estimate ampliconic gene family copy numbers. We extracted DNA from mouse liver using a Qiagen DNeasy kit and sent samples to Novogene (Novogene Corporation Inc., Sacramento, California) for library preparation and sequencing using Illumina HiSeq paired-end 150bp. Libraries were prepared and sequenced twice to increase unique coverage. We trimmed raw reads with Trimmomatic version 0.39 (Bolger et al. 2014). We mapped reads to the mouse reference genome build GRCm38 using bwa mem version 0.7.17 (Li and Durbin 2009) and used picard version 2.18.29 to fix mates and mark duplicates (Picard Toolkit). Data from the two sequencing runs were then merged for each sample.

To identify paralogs of ampliconic gene families, we extracted known X (*Slx, Slxl1, Sstx*), Y (*Sly, Ssty1, Ssty2*), and autosomal (*Speer*, and *α-takusan*) ampliconic gene sequences from the mouse reference GRCm38 using Ensembl annotation version 102 (Yates et al. 2019). We used the predicted gene *Gm5926* for *Sstx* because *Sstx* was not annotated in this version of Ensembl. For the autosomal gene families, we used the longest annotated genes in the gene family (*α*7-*takusan* and *Speer4f2*). We performed Ensembl BLAT searches with these sequences against the GRCm38 mouse reference, allowing up to 1000 hits. We then extracted all BLAT hits with greater than or equal to 97% sequence identity and an e-value of 0.0 and considered these filtered BLAT hits to be gene family paralogs for downstream copy number estimation.

We estimated copy numbers using a relative coverage approach similar to (Morgan and Pardo-Manuel De Villena 2017) and AmpliCoNE (Vegesna et al. 2020). For the relative coverage approach, we used Mosdepth v0.3.2 (Pedersen and Quinlan 2017) to estimate coverage across paralogous regions and divided this sum by half the genome-wide average coverage to account for hemizygosity of the sex chromosomes in males.

AmpliCoNE also estimates copy number based on relative coverage, while also controlling for GC content and only using informative regions based on repeat masking and mappability. AmpliCoNE was developed for estimating copy number on the assembly and annotation of the human Y, so we made some modifications to allow AmpliCoNE to work with the mouse sex chromosomes (Larson et al. 2021; https://github.com/ekopania/modified-AmpliCoNE). Specifically, we replaced AmpliCoNE’s method for identifying informative sites with an approach more suitable for the mouse assembly. For each ampliconic gene family, we extracted all k-mers of length 101bp from the sequence of one gene representing the ampliconic family and mapped these back to the mouse reference genome using Bowtie2 and allowing up to 500 multiple mapping hits. For each gene, we identified the most frequent number of times (*m*) k-mers mapped to the mouse genome and kept only k-mers that mapped *m* times. We identified all locations where these k-mers mapped with 2 or fewer mismatches. We considered the start locations of these k-mer mapping hits to be “informative sites.”

A small amount of autosomal material (∼0.1%) is expected to have introgressed along with the Y chromosome in our backcross experiments. To test this theoretical expectation and identify regions of introgression, we mapped whole genome sequence data from Y-introgression strains to both parental genomes using bwa mem v0.7.17-r1188 (Li and Durbin 2009) and called variants with GATK HaplotypeCaller v4.2.2.0. We then counted the number of variants in 100kb windows across the autosomes and identified regions where the number of variants when mapped to the maternal parent (autosomal background) genome exceeded the number of variants when mapped to the paternal parent (Y-introgression) genome. We repeated this analysis using whole genome sequence data from PWK and LEWES samples in our mouse colony. We excluded regions that had more variants when mapped to the opposite strain than when mapped to the same strain, as these are likely regions where genotype calls are unreliable due to assembly issues. After excluding these regions, 100kb windows with at least two more variants when mapped to the maternal parent compared to the paternal parent were considered introgressed in Y-introgression strains, reflecting the 95th percentile of differences in the number of variants within a window.

### Reproductive phenotypes

We phenotyped unmated male mice that were weaned at 21 days post-partum (dpp) into same-sex sibling groups and housed individually starting at 45 dpp to minimize effects of social dominance. Phenotype data were collected from at least six indivduals for each cross type; sample sizes for each phenotype and cross type are in Table 1. We weighed paired testes and paired seminal vesicles and calculated their mass relative to body weight. We compared offspring sex ratios from Y-introgression mice by recording the number of offspring of each sex at weaning. We then tested for a significant difference from an even sex ratio using a Pearson’s chi-squared test in R, and did a power analysis for this chi-squared test using the pwr.chisq.test function in the pwr package in R.

**Table 1.** Reproductive phenotypes for experimental XY mismatch mice and controls. Median values are presented +/1 standard error. Sample sizes are in parentheses. For sperm morphology parameters (bounding height, bounding width, area, perimeter, difference from median [a measure of the variability of nuclear shapes within the sample]), sample sizes indicate the number of sperm heads observed, and variance is depicted in violin plots (Figure 3; Supplemental Material, Figure S5). Gray boxes indicate significant differences (FDR-corrected Wilcoxon rank sum test P < 0.05) between XY mismatch cross types and control cross types with the same autosomal background. (‡) Indicates phenotypes with significant differences (FDR-corrected pairwise Wilcoxon rank sum test P < 0.05) between *mus*×*dom* F1 hybrids and both parental subspecies (*mus* and *dom*). (*) Indicates phenotypes with significant differences (FDR-correct pairwise Wilcoxon rank sum test P < 0.05) between *dom*×*mus* F1 hybrids and both parental subspecies (*mus* and *dom*). Testes and seminal vesicle weights are both paired. SV = seminal vesicle

| Phenotype | Hybrid F1 XY Match |  |  |  | Nonhybrid XY Mismatch |  |  |  |
| --- | --- | --- | --- | --- | --- | --- | --- | --- |
|  | *dom* × *mus* | *dom* × *mus*<sup>domY</sup> | *mus* × *dom* | *mus* × *dom*<sup>musY</sup> | *mus* | *mus*<sup>domY</sup> | *dom* | *dom*<sup>musY</sup> |
| Body mass (g) | 20 +/- 0.3 (24) | 19.6 +/- 0.3 (7) | 17.9 +/- 0.4 (24) | 18 +/- 0.4 (12) | 19 +/- 0.4 (23) | 18.4 +/- 0.3 (47) | 18 +/- 0.2 (67) | 19 +/- 0.5 (21) |
| Testes mass (mg)‡ | 186.4 +/- 3 (24) | 200.7 +/- 2 (6) | 123.9 +/- 2 (23) | 125.6 +/- 3 (12) | 193.2 +/- 5 (23) | 172.7 +/- 2 (47) | 209.1 +/- 3 (67) | 189.3 +/- 6 (21) |
| Relative testes mass (mg/g)*‡ | 9.1 +/- 0.1 (24) | 10.4 +/- 0.2 (6) | 7.2 +/- 0.1 (23) | 6.9 +/- 0.1 (12) | 10.2 +/- 0.2 (23) | 9.2 +/- 0.1 (47) | 11.7 +/- 0.1 (67) | 10.1 +/- 0.2 (21) |
| Relative SV mass (mg/g) | 6.6 +/- 0.2 (23) | 7.3 +/- 0.6 (6) | 5.2 +/- 0.3 (24) | 5.3 +/- 0.3 (12) | 6 +/- 0.3 (23) | 6.7 +/- 0.2 (47) | 5.2 +/- 0.2 (65) | 5.9 +/- 0.3 (21) |
| Bounding height‡* | 8.39 (1583) | 8.14 (650) | 7.46 (870) | 7.52 (847) | 8.21 (391) | 8.02 (401) | 8.11 (467) | 8.23 (443) |
| Bounding width‡* | 5.58 (1583) | 5.07 (650) | 4.02 (870) | 4.09 (847) | 5.02 (391) | 4.87 (401) | 4.9 (467) | 5.11 (443) |
| Area‡* | 24.5 (1583) | 21.6 (650) | 20.1 (870) | 20.1 (847) | 22.1 (391) | 20 (401) | 21.3 (467) | 20.4 (443) |
| Perimeter‡* | 23.8 (1583) | 22.7 (650) | 19.8 (870) | 20.2 (847) | 22.7 (391) | 21.9 (401) | 22.3 (467) | 23 (443) |
| Difference from median‡* | 6.22 (1583) | 8.67 (650) | 8.22 (870) | 10.8 (847) | 8.88 (391) | 5.77 (401) | 5.86 (467) | 6.72 (443) |

To quantify sperm morphology, we extracted sperm from each cross type from cauda epididymides diced in 1mL Dulbecco’s PBS (Sigma) and incubated at 37°C for 10 minutes. Sperm were fixed in 2% PFA, then dropped onto a slide with DAPI solution to stain the sperm nuclei. We imaged greater than 400 nuclei per genotype and analyzed the images using the Nuclear Morphology Analysis software (Skinner et al. 2019). We used two microscopes but performed clustering analysis on combined nuclei imaged from both microscopes to ensure that nuclei imaged on one scope were not clustering separately from those taken on the other microscope (Supplemental Material, Figure S1). The Nuclear Morphology Analysis software uses a Canny edge detection algorithm to detect objects (nuclei) within images, orients and aligns the nuclei, and uses a modification of the Zahn-Roskies transformation of the nucleus outlines to automatically detect landmarks. The software estimates area, perimeter, bounding height, bounding width, regularity, difference from median, and a consensus shape of the nuclei for each genotype. We tested for significant differences among cross types for each of these parameters using a Wilcoxon rank sum test in R. Using this automated morphology analysis software, we were able to analyze 5652 nuclei and detect subtle but significant differences that may not be measurable by eye or qualitative analysis.

### Testis sorting and RNA sequencing

We collected testes from mice immediately following euthanization and isolated cells at different stages of spermatogenesis using Fluorescence-Activated Cell Sorting (FACS; Getun et al. 2011). The full FACS protocol is available on GitHub (https://github.com/goodest-goodlab/good-protocols/tree/main/protocols/FACS). Briefly, we decapsulated testes and washed them twice with 1mg/mL collagenase (Worthington Biochemical), 0.004mg/mL DNase I (Qiagen), and GBSS (Sigma), followed by disassociation with 1mg/mL trypsin (Worthington Biochemical) and 0.004mg/mL DNase I. We then inactivated trypsin with 0.16mg/mL fetal calf serum (Sigma). For each wash and disassociation step, we incubated and agitated samples at 33°C for 15 minutes on a SciGene Model 700 Microarray Oven at approximately 10rpm. We stained cells with 0.36mg/mL Hoechst 33324 (Invitrogen) and 0.002mg/mL propidium iodide and filtered with a 40μm cell filter. For Hybrid F1 XY Match, we sorted using a FACSAria Fusion flow cytometer, and for Nonhybrid XY Mismatch we sorted cells using a FACSAria IIu cell sorter (BD Biosciences), both at the UM Center for Environmental Health Sciences Fluorescence Cytometry Core. We periodically added 0.004mg/mL DNase I as needed during sorting to prevent DNA clumps from clogging the sorter. We sorted cells into 15μL beta-mercaptoethanol (Sigma) per 1mL of RLT lysis buffer (Qiagen) and kept samples on ice whenever they were not in the incubator or the cell sorter. We performed cell sorting on four individuals of each cross type and focused on two cell populations: early meiotic spermatocytes (leptotene/zygotene) and postmeiotic round spermatids. We extracted RNA using the Qiagen RNeasy Blood and Tissue Kit and checked RNA integrity with a TapeStation 2200 (Agilent). Only two samples had RNA integrity numbers (RIN) less than 8 (RIN = 7 and 7.1; Supplemental Material, Table S1). We prepared RNAseq libraries using the KAPA mRNA hyperprep kit and sequenced samples with Novogene (Illumina NovaSeq6000 PE 150). Samples were prepared and sequenced together, but Hybrid F1 XY Match mice and Nonhybrid XY Mismatch mice were sorted on different FACS machines, so to minimize experimental batch effects we analyzed these two experiments separately unless otherwise noted.

### Gene expression analyses

We performed gene expression analyses on FACS expression data representing two cell populations: early meiosis (leptotene-zygotene, hereafter “early”) and postmeiosis (round spermatids, hereafter “late”). For the early cell type, a few samples did not group with others of the same cross type in multidimensional scaling (MDS) plots (Supplemental Material, Figure S2). These samples were likely contaminated with other cell types based on their relative expression levels of cell-type marker genes from *Mus musculus* testes single-cell RNAseq experiments (Supplemental Material, Figure S3; Green et al. 2018; Hunnicutt et al. 2021), and were therefore removed from expression analyses. Because sex chromosome ampliconic genes are primarily expressed in late spermatogenesis (Mueller et al. 2013; Larson et al. 2018a), and disrupted sex chromosome expression in hybrid males primarily occurs after the early cell type stage (Larson et al. 2017), we focus on data from the late cell type in the main text and report results from the early cell type in the Supplemental Material.

We performed gene expression analyses using mice from both our Hybrid F1 XY Match and Nonhybrid XY Mismatch experiments, and reanalyzed expression data from (Larson et al. 2017), which generated spermatogenesis cell-type enriched gene expression data from the same F1 hybrid crosses (PWK^♀^ × LEWES^♂^ and LEWES^♀^ × PWK^♂^) and intrasubspecific F1 crosses (CZECHII^♀^ × PWK^♂^ and WSB^♀^ × LEWES^♂^) used in this study.

We trimmed RNAseq reads using trimmomatic v0.39 (Bolger et al. 2014). One sample (PP.LL30.7MLZ) had about an order of magnitude more reads than any other sample (> 900 million raw reads), so we downsampled to the mean number of reads after trimming using fastq-sample version 0.8.3 and verified that reads were properly paired after downsampling using fastq_pair (Edwards and Edwards 2019). We quantified reads using a kmer-based quasi-mapping approach implemented in salmon v1.4.0 (Patro et al. 2017) and a salmon index based on the mouse reference transcriptome version GRCm38. We then converted from transcript-level counts to gene-level counts using the R packages tximport 1.14.2 and EnsDb.Mmusculus.v79. We used EdgeR version 3.32.1 to normalize expression data. First, we filtered out genes with low expression by only including genes that had an FPKM > 1 in at least 4 samples. Then, we normalized expression data following the recommendations in the tximport documentation.

We quantified expression levels of ampliconic gene families by calculating transcripts per million (TPM) for each gene separately then summing TPM values for all paralogs of a gene family (≥97% sequence identity). We used linear mixed-effect models to test if gene family expression level was significantly associated with copy number for *Slx, Slxl1, Sly, Ssty1, Ssty2*, and *α*-*takusan*. We compared disrupted expression levels on the autosomes, X chromosome, and Y chromosome by subtracting normalized FPKM values in control mice from normalized FPKM values in XY mismatch mice and control mice for every gene (Good et al. 2010). We then used a Mann-Whitney U test to compare the distribution of normalized FPKM differences among the chromosome types. To identify Differentially Expressed (DE) genes between cross types, we used the likelihood ratio test approach with false-discovery rate (FDR) correction in EdgeR and visualized overlaps in DE genes among cross types using the R package UpSetR (Conway et al. 2017). We removed DE genes in autosomal regions we identified as putatively introgressed, because these genes may be DE due to introgressed autosomal variants rather than incompatibilities resulting from mismatching sex chromosomes. For ampliconic genes with high sequence similarity, some reads are expected to map multiply but will only be assigned to one member of the ampliconic gene family. Therefore, individual genes within gene families may sometimes be identified as DE, even though their paralogs are not, due to differences in read assignment across paralogs.

We further investigated genome-wide expression differences among cross types using weighted correlation network analyses (WGCNA; Langfelder and Horvath 2008). We identified correlated expression modules significantly associated with different cross types using a linear model and Tukey’s honest significant difference (HSD) test. We used R version 4.0.3 for all statistical tests and to implement all R packages (R Core Team).

## Results

### Copy Number Imbalance in Y-introgression Mice

We first estimated ampliconic gene family copy numbers in wild mice, wild-derived inbred strains, and Y-introgression mice using whole genome sequencing. The samples that we sequenced had genome-wide average coverages of 10-15×, and samples with publicly available data all had coverage >5×. We found that *musculus* tended to have higher *Slx* and *Sly* copy numbers than *domesticus* (median *Slx* copy number in *musculus*: 62, in *domesticus*: 17, FDR-corrected Wilcoxon rank sum P < 0.01; median *Sly* copy number in *musculus*: 226, in *domesticus*: 109, FDR-corrected Wilcoxon rank sum P < 0.01), qualitatively consistent with previous studies (Ellis et al. 2011; Case et al. 2015; Morgan and Pardo-Manuel De Villena 2017; Figure 2A). *Slxl1* copy numbers also tended to be higher in *musculus*, but there was high copy number variation for this gene family in *domesticus* with some samples reaching copy numbers as high as those found in *musculus* (median *Slxl1* copy number in *musculus*: 37, in *domesticus*: 31, FDR-corrected Wilcoxon rank sum P < 0.01; Figure 2B). *Slx, Slxl1*, and *Sly* copy numbers for wild-derived inbred strains were representative of those found in wild mice (Figures 2A and 2B; Supplemental Material, Table S3), consistent with previous results (Larson et al. 2021). Our Y-introgression mice retained copy numbers similar to those of pure strains with the same X and Y chromosome genotypes, so they had *Slx*-*Sly* and *Slxl1*-*Sly* dosage imbalance similar to that expected in natural hybrids (Figures 2A and 2B; Supplemental Material, Table S3).

**Figure 2.**
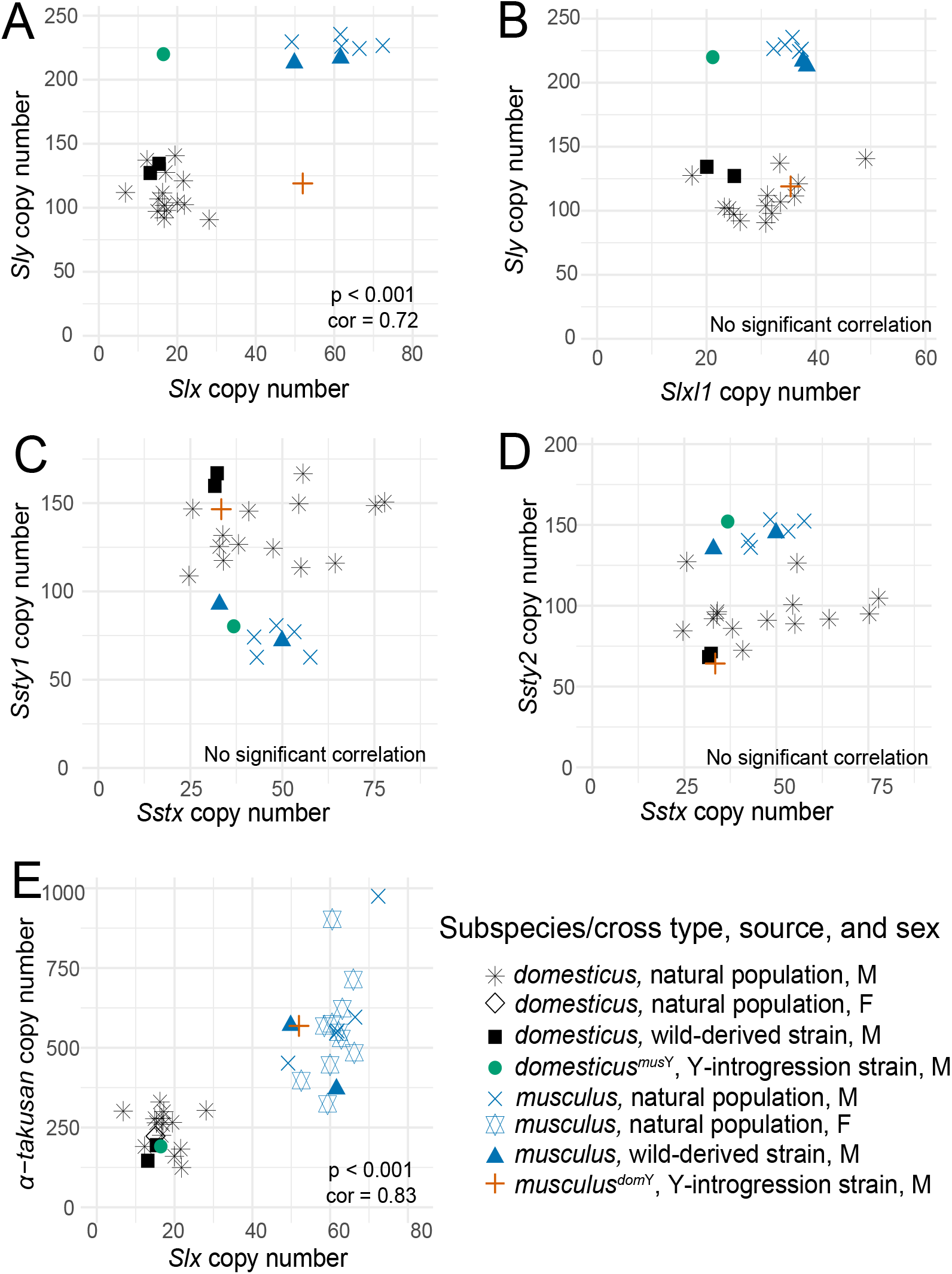
Copy number estimates for ampliconic gene families in wild mice, wild-derived inbred strains, and Y-introgression strains. Copy number was estimated using a 97% identity cutoff for paralogs. (A-D) show copy numbers in male mice, with Y chromosome genes on the y-axis and their X chromosome homologs on the x-axis. (E) includes both males and females and shows haploid copy number for the autosomal gene family *α*-*takusan* on the y-axis and haploid copy number for the X-linked family *Slx* on the x-axis. Note that (A) and (B) show the same information on the y-axis and (C) and (D) show the same information on the x-axis to compare copy numbers for ampliconic gene families that have two different homologous gene families on the opposite sex chromosome. Correlations and p-values are based on a Pearson’s correlation test. P-values were FDR-corrected for multiple tests.

**Figure 3.**
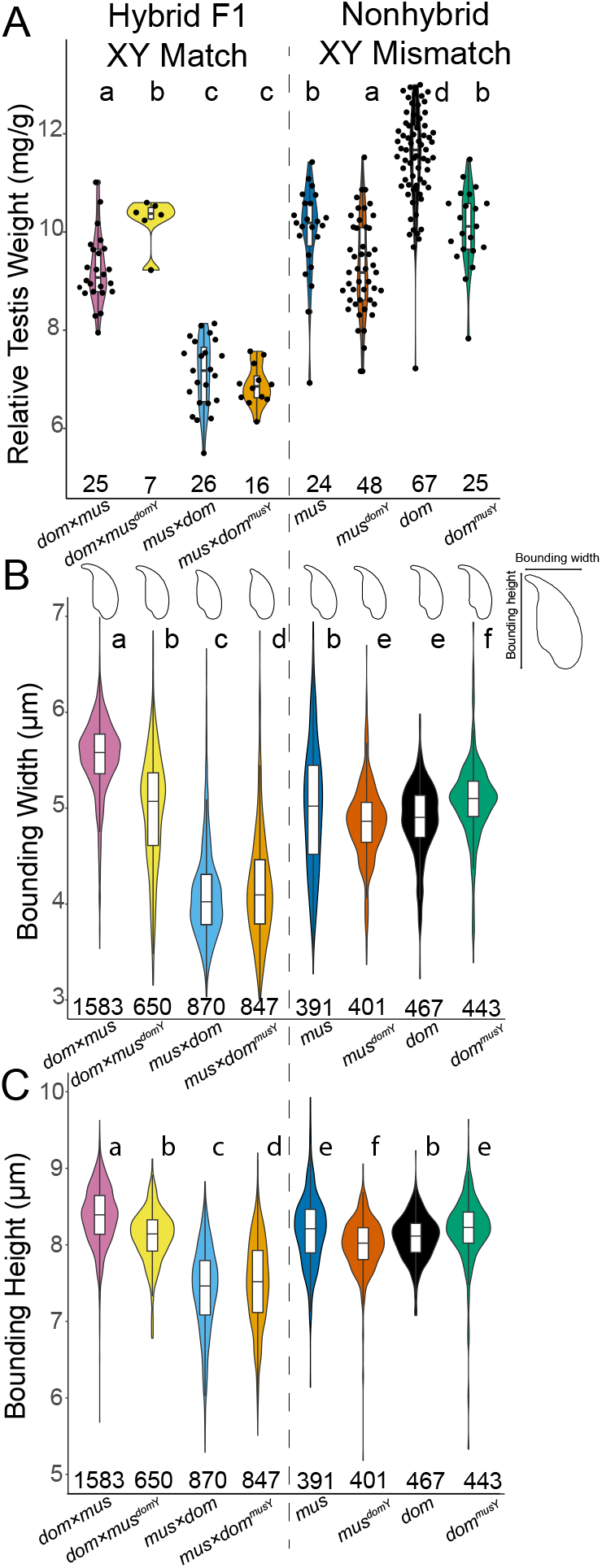
(A) Relative testes mass (mg/g), (B) sperm nucleus bounding width (μm), and (C) sperm nucleus bounding height (μm) by cross type. Letters above each violin plot indicate significant differences (FDRcorrected P < 0.05) based on a Welch’s t-test (relative testes mass) or Wilcoxon rank-sum test (bounding width and height). Sample size for each cross type is indicated below each violin plot. Bounding width and height sample sizes indicate the number of sperm nuclei observed. Representative sperm nuclei morphologies for each cross type are depicted above each violin plot in (B).

Additional ampliconic gene families showed copy number differences between *musculus* and *domesticus* that were also represented in our Y-introgression mice. *Sstx* had similar copy numbers in *musculus* and *domesticus*, but its two Y-linked homologs showed differences between subspecies, with *Ssty1* having more copies in *domesticus* and *Ssty2* having more copies in *musculus* (median *Sstx* copy number in *musculus*: 48, in *domesticus*: 39, FDR-corrected Wilcoxon rank sum P = 0.57; median *Ssty1* copy number in *musculus*: 74, in *domesticus*: 139, FDR-corrected Wilcoxon rank sum P < 0.01; median *Ssty2* copy number in *musculus*: 145, in *domesticus*: 92, FDR-corrected Wilcoxon rank sum P < 0.01; Figure 2C, 2D).

We also estimated copy number for *α*-*takusan* and *Speer*, two autosomal ampliconic gene families thought to be regulated by sex chromosome ampliconic genes (Moretti et al. 2020). In both males and females, *α*-*takusan* showed a high correlation in copy number with *Slx* (r = 0.95; Pearson’s correlation P < 0.001), suggesting that it was co-amplified with the *Slx* gene family (Figure 2E). Note that correlation tests were performed without phylogenetic correction, because we wanted to test if gene families were co-amplified regardless of whether this was a result of shared evolutionary history. *Speer* copy number was more difficult to estimate using our approaches due to lower sequence similarity among *Speer* paralogs compared to other ampliconic gene families, but our estimates suggested that *Speer* may also have higher copy number in *musculus* relative to *domesticus* (Supplemental Material, Table S3). To verify our computational copy number estimates, we also performed digital droplet PCR (ddPCR) on a subset of *dom* samples using the *Slxl1* primers from (Kruger et al. 2019). We found 15 *Slxl1* copies with ddPCR, consistent with findings in (Kruger et al. 2019). While our computational estimates are higher than this, we found similar results if we imposed a stricter cutoff for considering genes paralogs (98-99% sequence identity), likely reflecting a high specificity of the primers we used. We also found similar results using a different computational approach based on relative coverage (Supplemental Material, Table S3; Larson et al. 2021).

### Residual Autosomal Introgression in Y-introgression Strains

We identified putative introgressed regions by mapping samples to both subspecies reference genomes, dividing the reference genome autosomal regions into 24,639 100kb windows, and identifying SNPs in these windows. We found evidence for introgression in 105 windows in *domesticus*^*mus*Y^, and 33 windows in *musculus*^*dom*Y^, representing 0.43% and 0.13% of the autosomal windows that passed filtering, respectively (Supplemental Material, Table S4). Thus, the *domesticus*^*mus*Y^ strain had approximately four times more introgression than the theoretical expectation of 0.1% based on the number of backcross generations. The relatively large difference in percentages of introgression between the strains was primarily due to an ∼7.6 Mbp introgressed region on chromosome 2 in *domesticus*^*mus*Y^ (Supplemental Material, Figure S4). This large introgressed region had an average difference of 958 SNPs, in contrast to the median difference of eight SNPs across all other putatively introgressed autosomal regions. Thus, the introgressed region on chromosome 2 in the *domesticus*^*mus*Y^ strain likely represents the only large track of autosomal introgression, with some evidence for additional, smaller amounts of introgression throughout the autosomes in both reciprocal Y-introgression strains.

Some of the putatively introgressed regions we identified may be prone to introgression more generally. The large area on chromosome 2 overlapped with a region with evidence for introgression from *musculus* into the *domesticus* wild-derived inbred strains STRA and STRB (Mukaj et al. 2020). We used the Mouse Phylogeny Viewer (Yang et al. 2011) to identify an additional nine mouse inbred strains with introgression from *musculus* into a *domesticus* background in this region (Supplemental Material, Figure S4C). In one area of the mouse hybrid zone, a SNP contained within this introgressed region showed evidence for excess of the *musculus* allele in mice with primarily *domesticus* backgrounds, suggesting that introgression of this region from *musculus* into *domesticus* may have occurred in wild populations (Teeter et al. 2010). This region is also adjacent to *R2d2*, a copy number variant in mice that shows transmission ratio distortion in females heterozygous for the high copy number *R2d2* drive allele (Didion et al. 2016). We also identified 5 different 100kb windows near each other on chromosome 14 with evidence for introgression in *musculus*^*dom*Y^ mice that overlap with a region in the *musculus* wild-derived strain PWD with evidence for introgression from *domesticus* (41.3-41.4Mb, 41.8-41.9Mb, 42.2-42.3Mb, 42.3-43.4Mb, and 44.2-44.3Mb; Mukaj et al. 2020).

### XY Mismatch Contributed to Some Male Sterility Phenotypes

We next asked if XY mismatch was associated with male sterility phenotypes (Table 1). For Hybrid F1 XY Match, where we compared hybrid mice both with and without sex chromosome mismatch, hybrids with a *musculus*^♀^ × *domesticus*^♂^ background had lower relative testes mass than hybrids with the reciprocal *domesticus*^♀^ × *musculus*^♂^ background regardless of whether they had XY mismatch or not (Figure 3A). These results were consistent with previous studies showing more severe hybrid sterility in the *musculus*^♀^ × *domesticus*^♂^ direction of this cross (Good et al. 2008b; Good et al. 2010; Campbell et al. 2012; Larson et al. 2017). Although *domesticus*^♀^ × *musculus*^♂^ showed much less severe sterility phenotypes than the reciprocal F1 hybrid, we still considered these mice to be potentially subfertile because their relative testes mass and sperm morphology parameters were significantly different from those of either pure *dom* or pure *mus* (Figure 3, Table 1), and even subtle reductions in fertility may be important in nature, where sperm competition is high for house mice (Dean et al. 2006). For Hybrid F1 XY Match mice, *dom*×*mus*^*dom*Y^ mice had higher relative testis mass than *dom*×*mus* mice, suggesting that XY match partially rescued relative testes mass in some mice with a hybrid autosomal background (Figure 3A). In the reciprocal direction, however, XY match had no significant effect on relative testes mass (Figure 3A). For Nonhybrid XY Mismatch, we found that mice with XY mismatch had reduced relative testis mass compared to control mice with the same nonhybrid X and autosomal background (Figure 3A). In summary, we found little effect of XY mismatch on testis mass in the most sterile F1 cross (*musculus*^♀^ × *domesticus*^♂^), where sterility is therefore likely due to X-autosomal or autosomal-autosomal incompatibilities (Campbell and Nachman 2014). However, in the reciprocal and more fertile F1 direction XY mismatch seemed to have an important effect on testis mass. Furthermore, in the absence of any autosomal or X-autosomal incompatibilities, XY mismatch resulted in slightly but significantly decreased relative testis mass.

We saw severe sperm head abnormalities in our Hybrid F1 XY Match crosses with a *musculus*^♀^ × *domesticus*^♂^ background (*mus*×*dom* and *mus*×*dom*^*mus*Y^). Sperm from both these cross types had significantly lower bounding height and bounding width compared to all other cross types (FDR-corrected Wilcoxon rank sum P << 0.0001; Table 1), largely due to their shortened hook and consistent with hybrid sterility in this direction of the cross (Figure 3B, 3C). This was also consistent with previous manual (categorical) observations of abnormal sperm head morphology in this cross type in other studies (Good et al. 2008a; Campbell and Nachman 2014; Larson et al. 2017; Larson et al. 2018b). The reciprocal *dom*×*mus* F1 hybrids had sperm with higher bounding height and bounding width compared to sperm from all other cross types, including the reference subspecies (FDR-corrected Wilcoxon rank sum P < 0.01; Table 1; Figure 3B, 3C). This direction of the cross is generally considered more fertile but sometimes shows reduced fertility compared to nonhybrid mice (Larson et al. 2018b). It is possible that the larger overall size of these sperm may reflect abnormal nuclear packaging and could contribute to reduced fertility in *domesticus*^♀^ × *musculus*^♂^ F1 mice. When comparing XY match mice to F1 hybrids with abnormally small sperm heads, *mus*×*dom*^*mus*Y^ mice had significantly higher bounding width and bounding height than *mus*×*dom* mice (FDR-corrected Wilcoxon rank sum P < 0.01; Table 1; Figure 3B, 3C). These results suggest that XY match rescued some of the aberrant sperm head morphology associated with hybrid sterility in *musculus*^♀^ × *domesticus*^♂^ F1s, but the effects of XY match rescue were subtle, consistent with previous observations (Campbell and Nachman 2014). In the reciprocal cross direction, *dom*×*mus*^*dom*Y^ had lower bounding width and bounding height than the abnormally large *dom*×*mus* sperm heads (FDR-corrected Wilcoxon rank sum P << 0.0001; Table 1; Figure 3B, 3C), so XY match rescued some of the oversized sperm head morphology we observed in *dom*×*mus*. In Nonhybrid XY Mismatch, we observed subtle effects of XY mismatch consistent with our Hybrid F1 XY Match observations. Sperm from *mus*^*dom*Y^ mice had slightly lower bounding height and bounding width compared to sperm from *mus* (FDR-corrected Wilcoxon rank sum P < 0.01; Table 1; Figure 3B, 3C), consistent with lower bounding height and bounding width in sperm from *mus*×*dom* mice that also had a *mus* X chromosome and *dom* Y chromosome. However, *mus*^*dom*Y^ sperm were more similar in size to *mus* sperm than *mus*×*dom* sperm and qualitatively had a hook morphology more similar to that of fertile *mus* than sterile *mus*×*dom* mice, so the contribution of XY mismatch to sperm head morphology is small compared to the effect of X-autosomal interactions. In the reciprocal direction, *dom*^*mus*Y^ mice had sperm with higher bounding height and bounding width compared to sperm from *dom* mice (FDR-corrected Wilcoxon rank sum P << 0.0001; Table 1; Figure 3B, 3C), consistent with the higher bounding height and bounding width in *dom*×*mus* hybrids. Sperm from *dom*^*mus*Y^ mice also had smaller areas (FDR-corrected Wilcoxon rank sum P << 0.0001; Table 1; Supplemental Material, Figure S5), so the larger bounding height and bounding width are primarily the result of a slightly elongated hook rather than an overall increase in the sperm head size. Other sperm head morphology parameters, including area, perimeter, and differences from median, showed similar subtle differences or no differences among cross types (Table 1; Supplemental Material, Figures S1 and S5).

Genetic manipulation studies have shown offspring sex ratio skews under *Slxl1*-*Sly* dosage imbalance, contributing to evidence for *Slxl1*-*Sly* intragenomic conflict. Male mice with an excess of *Sly* relative to *Slxl1* produce more male offspring, while mice with an excess of *Slxl1* produce more female offspring (Cocquet et al. 2012; Kruger et al. 2019) due to reduced motility of Y-bearing sperm (Rathje et al. 2019). We asked if more subtle imbalances in relative copy numbers expected in natural hybrid mice also result in sex ratio skews and did not see a significant difference from a 50:50 sex ratio for offspring of XY mismatch mice (Supplemental Material, Table S5). A more extreme dosage imbalance than that seen in our XY mismatch experimental mice (and in natural hybrids) is probably required to produce a large sex ratio skew. However, it is important to note that we had very little power to detect differences in sex ratio, with type II error probabilities over 0.8 (Supplemental Material, Table S5).

### *Slx/Slxl1*-*Sly* Dosage Imbalance Did Not Lead to Ampliconic Gene Family Overexpression

Copy number imbalance of *Slx* and *Slxl1* relative to *Sly* is thought to disrupt expression of these gene families in late spermatogenesis, with particularly strong evidence for *Slx* and *Slxl1* overexpression when *Sly* is knocked down (Cocquet et al. 2009; Cocquet et al. 2012) and *Slxl1* overexpression when *Slx* and *Slxl1* are duplicated (Kruger et al. 2019). *Slx, Slxl1*, and *Sly* appear to be involved in the regulation of sex chromatin which impacts the regulation of many genes during late spermatogenesis (Kruger et al. 2019). Therefore, we predicted that their misregulation may disrupt the expression of additional genes, including additional Y-linked ampliconic gene families *Ssty1*/2 and the autosomal ampliconic gene family *α*-*takusan* (Larson et al. 2017; Moretti et al. 2020). To test if *Slx, Slxl1, Sly, Ssty1, Ssty2*, and *α*-*takusan* expression was disrupted under less extreme copy number differences in hybrid mice, we compared ampliconic gene family expression levels in round spermatids among cross types. We did not directly quantify copy number for the mice that were FACS sorted, so we used our previous copy number estimates from pure strains sharing the same sex chromosomes as our experimental mice (Larson et al. 2021). For all six gene families, expression level was significantly associated with copy number based on a linear mixed-effects model with experiment as a random effect to control for batch effects (FDR-corrected P < 0.05; Figure 4). However, for *Slxl1*, this association was negative, suggesting that copy number was not the primary determinant of *Slxl1* expression. This is interesting given that we found high overlap in the range of *Slxl1* copy numbers in naturally occurring *musculus* and *domesticus* (Figure 2B), and the previous demonstration that *Slxl1* plays a more direct role in sex ratio bias than *Slx* (Kruger et al. 2019). We then tested if XY mismatch had a significant effect on expression level using a linear mixed-effects model with both copy number and presence of XY mismatch as fixed effects and experiment as a random effect. We used an ANOVA to compare this model to a null model with copy number as the only fixed effect and experiment as a random effect. For all six genes, XY mismatch was not significantly associated with ampliconic gene expression levels (FDR-corrected ANOVA P > 0.05). When we specified the direction of XY mismatch (i.e., *musculus* X and *domesticus* Y, the direction with an excess of *Slx* relative to *Sly*), only *Ssty2* expression was significantly associated with XY mismatch in this direction (FDR-corrected ANOVA P < 0.05).

**Figure 4.**
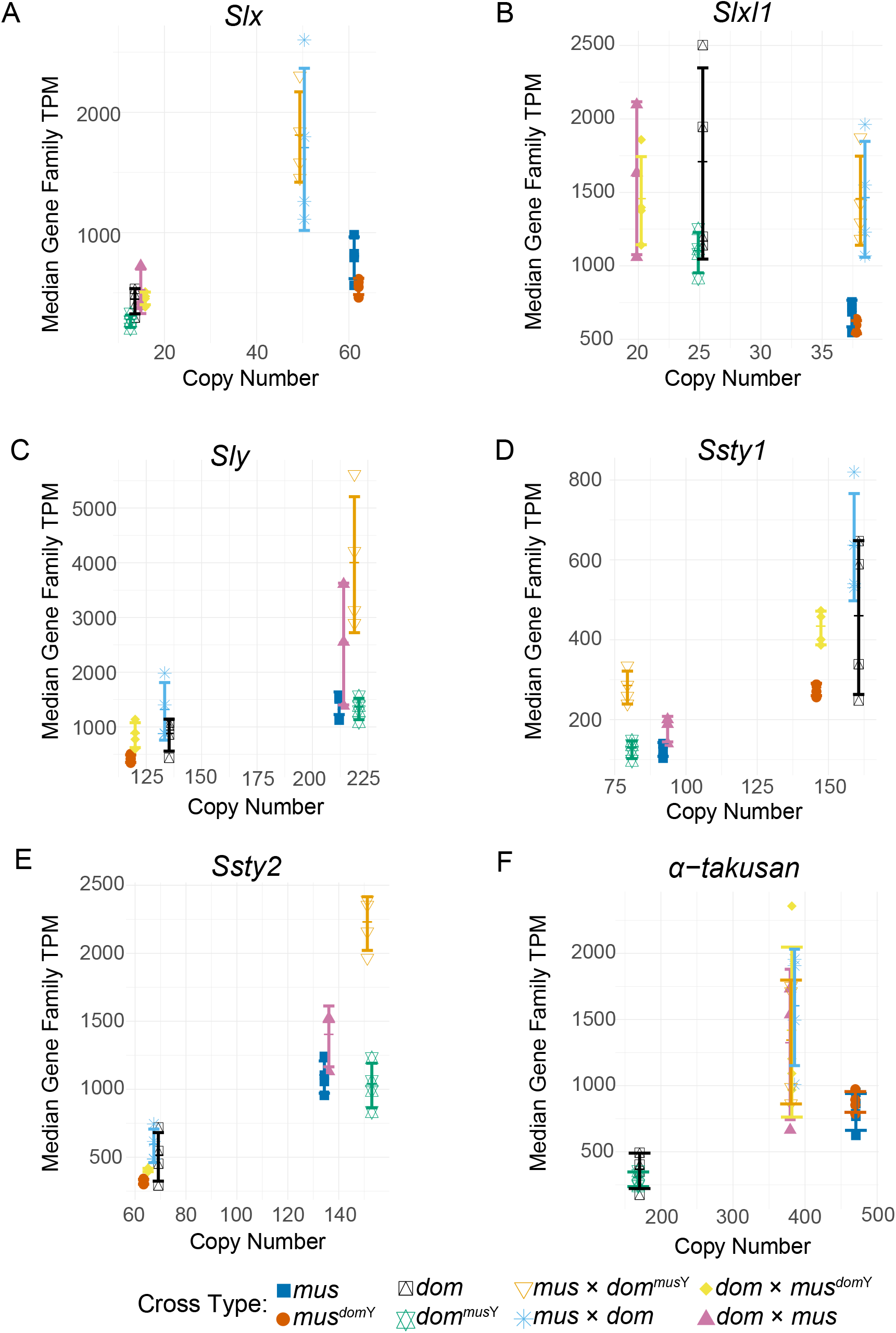
Normalized expression levels of *Slx* (A), *Slxl1* (B), *Sly* (C), *Ssty1* (D), *Ssty2* (E), and *α-takusan* (F) ampliconic gene families in different cross types plotted against their copy numbers. Copy number estimates are based on estimates from wild-derived strains used in experimental and control crosses (see Figure 2). Cross types with the same sex chromosome and therefore same copy number estimate are jittered slightly along the x-axis for clarity. Expression level was calculated by summing transcripts-per million (TPM) for each paralog of the gene family with at least 97% sequence identity to the ampliconic gene. Points represent values for individual samples, and lines indicate median and standard deviation for each cross type.

We also tested if X-autosomal background was significantly associated with expression levels using the same mixed-effects model approach. For *Slx, Slxl1, Sly, Ssty1*, and *Ssty2*, the sterile hybrid background (*musculus*^♀^ × *domesticus*^♂^) was significantly associated with expression levels after FDR-correction (*Slx* ANOVA P << 0.0001; *Slxl1* P < 0.001; *Sly* P = 0.01; *Ssty1* P < 0.001; *Ssty2* P = 0.001). We observed overexpression of *Slx, Slxl1, Sly, Ssty1*, and *Ssty2* relative to their copy numbers for mice with *musculus*^♀^ × *domesticus*^♂^ backgrounds (*mus*×*dom* and *mus*×*dom*^*mus*Y^; Figure 4A-E), consistent with previous studies showing that these hybrid mice exhibit widespread overexpression on the sex chromosomes (Good et al. 2010; Campbell et al. 2013; Larson et al. 2017). Both *mus*×*dom* and *mus*×*dom*^*mus*Y^ mice in our study overexpressed *Slx, Slxl1*, and *Sly* (Figure 4A, 4B, and 4C), suggesting that matching X and Y chromosomes from *musculus* did not rescue *Slx, Slxl1*, or *Sly* upregulation, and that the overexpression we observed likely results from X-autosomal incompatibilities that disrupt MSCI rather than *Slx*- or *Slxl1*-*Sly* dosage imbalance. Additionally, *mus*^*dom*Y^ mice from our Nonhybrid XY Mismatch also had a *musculus* X and *domesticus* Y, the same X and Y chromosome combination found in sterile hybrids that results in an excess of *Slx* and *Slxl1* copies relative to *Sly* copies. If *Slx*- or *Slxl1*-*Sly* dosage imbalance contributed to *Slx, Slxl1*, and *Sly* overexpression, we would expect *mus*^*dom*Y^ mice to have higher expression than *mus* controls. We observed the opposite effect, with *mus*^*dom*Y^ mice showing slightly lower *Slx, Slxl1*, and *Sly* expression levels (Figure 4A, 4B, and 4C). This result provides further evidence that postmeiotic *Slx, Slxl1*, and *Sly* overexpression in sterile F1 hybrids is unlikely to be primarily due to *Slx*- or *Slxl1*-*Sly* dosage imbalance, and that XY mismatch in the absence of autosomal mismatch is not sufficient to cause overexpression of *Slx, Slxl1*, and *Sly*.

Given that *Slx, Slxl1*, and *Sly* are thought to regulate the *α*-*takusan* ampliconic family, we predicted that *α*-*takusan* expression levels would also be associated with a *musculus*^♀^ × *domesticus*^♂^ background. Surprisingly, this association was not significant (ANOVA P = 0.40). Instead, we observed that *α*-*takusan* was overexpressed in all cross types with an F1 autosomal background regardless of cross direction (Figure 4F), and that expression was significantly associated with an F1 autosomal background (ANOVA P < 0.01). This suggests that *α*-*takusan* regulation likely involves autosomal loci in addition to SLX, SLXL1, SLY, SSTY1, and SSTY2 (Moretti et al. 2020).

Sex-linked ampliconic genes are primarily expressed during postmeiotic spermatogenesis, in mice and more generally across mammals (Cocquet et al. 2012; Mueller et al. 2013; Sin and Namekawa 2013). Our nonhybrid expression data supported this, with little to no expression of *Slx, Slxl1, Sly*, or *Ssty1*/2 in early meiotic cells in our *mus* and *dom* samples. However, we did detect some meiotic expression of *Slx, Slxl1, Sly*, and *Ssty2* in mice with hybrid autosomal backgrounds, and expression levels of these gene families in early meiosis was significantly associated with F1 autosomal background (ANOVA P < 0.05, Supplemental Material, Figure S6). X chromosome expression has been shown to be disrupted throughout spermatogenesis in F1 hybrids, although the effect was smaller during earlier spermatogenic stages (Larson et al. 2017). Our results suggest that disruption of early spermatogenesis regulatory networks may result in spurious expression of sex-linked ampliconic genes during early meiotic stages when they are normally silenced.

### XY Mismatch Was Not Associated with Sex Chromosome Overexpression in Sterile F1 Hybrids

Next we sought to differentiate if widespread postmeiotic overexpression in sterile hybrids was a direct result of sex chromosome mismatch, a continuation of disrupted meiotic sex chromosome inactivation (MSCI), or a combination of both (Larson et al. 2017; Larson et al. 2021). We first reanalyzed data from (Larson et al. 2017) and repeated their result showing sex chromosome upregulation in late spermatogenesis in sterile F1 hybrids (*mus*×*dom*, Figure 5A and 5D). We then tested if upregulation was due to XY mismatch by comparing relative expression levels in F1 hybrids to those in our Hybrid F1 XY Match mice, which had sex chromosomes from the same subspecies. If XY mismatch contributed to sex chromosome upregulation in sterile hybrids, we would expect to see some rescue from disrupted postmeiotic expression in these Hybrid F1 XY Match mice, with *mus*×*dom*^*mus*Y^ mice having lower expression on the X chromosome relative to *mus*×*dom* F1s. Contrary to this prediction, the X chromosome showed similar expression levels when comparing expression in these two cross types. Therefore, restoring matching sex chromosomes did not rescue expression levels on the *musculus* X chromosome from overexpression in hybrids (Figure 5B). We further tested the effects of sex chromosome mismatch using our Nonhybrid XY Mismatch mice, which had introgressed Y chromosomes on a nonhybrid autosomal background. If mismatch between a *musculus* X chromosome and *domesticus* Y chromosome was sufficient to induce postmeiotic sex chromosome overexpression, then we would expect to see higher X chromosome expression in *mus*^*dom*Y^ mice. Instead, we observed slight under expression on the X chromosome compared to the autosomes in *mus*^*dom*Y^ mice, confirming that sex chromosome mismatch does not cause X chromosome overexpression in late spermatogenesis (Figure 5C).

**Figure 5.**
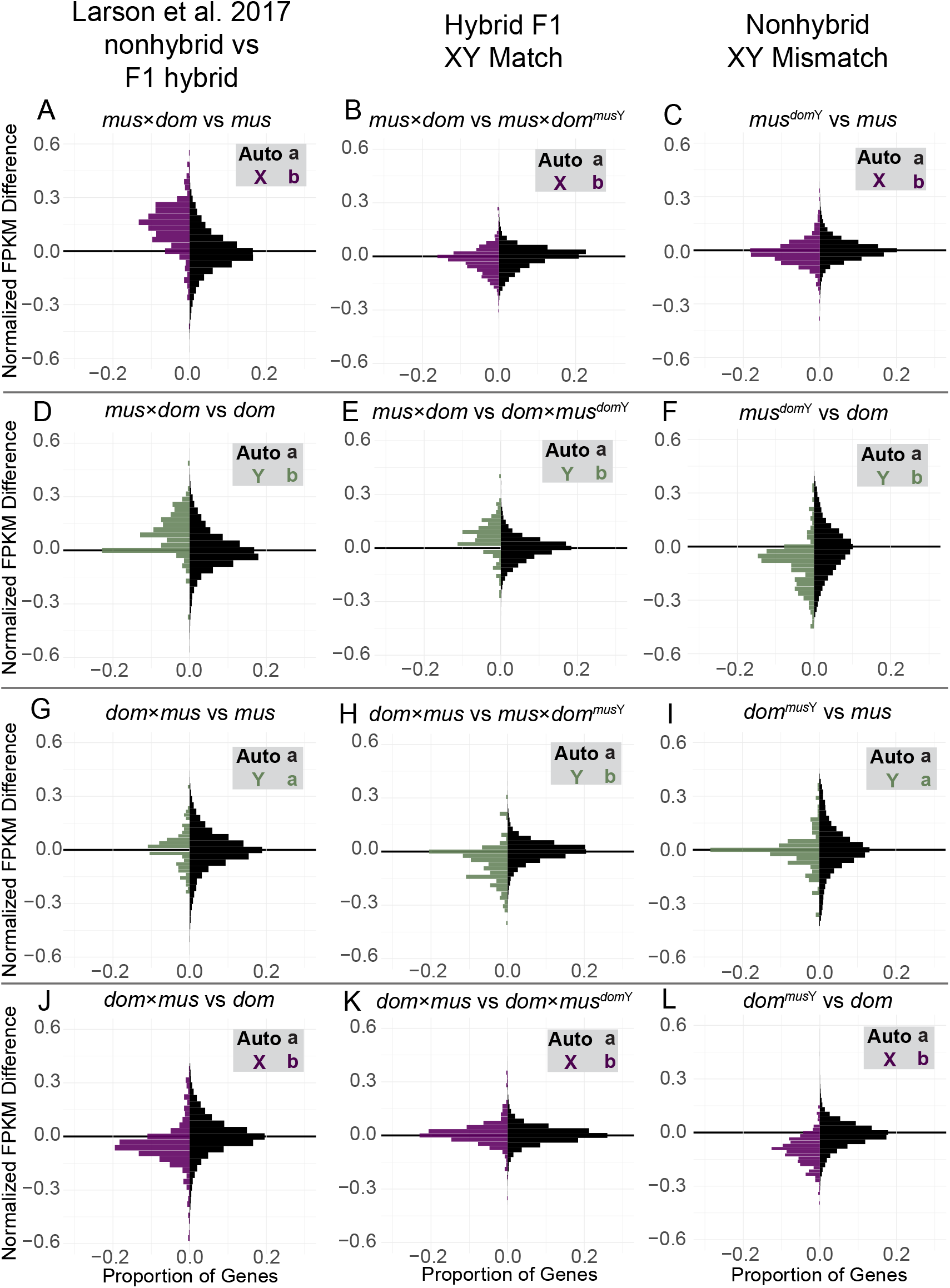
Histograms of relative expression levels between experimental cross types and control mice. (A-C) Contrasts that all have a *musculus* X chromosome, (D-F) contrasts with a *domesticus* Y chromosome (G-I) contrasts with a *musculus* Y chromosome, and (J-L) contrasts with a *domesticus* X chromosome. (A-F) represent sex chromosome mismatch present in sterile hybrids (*musculus* X and *domesticus* Y), while (G-L) represent sex chromosome mismatch present in more fertile hybrids (*domesticus* X and *musculus* Y). The first column (A, D, G, and J) shows data reanalyzed from (Larson et al. 2017). The second column (B, E, H, K) tests if gene expression levels are rescued when the sex chromosomes are matched but on a hybrid autosomal background (Hybrid F1 XY Match). The third column (C, F, I, L) tests for disrupted expression due to sex chromosome mismatch alone, on a nonhybrid autosomal background (Nonhybrid XY Mismatch). The y-axis shows the difference in normalized expression levels between the two cross types being compared. The x-axis shows the proportion of genes in each expression difference bin. Black bars represent the autosomes, purple bars represent the X chromosome, and green bars represent the Y chromosome. Letters indicate significant differences in median expression differences among the chromosome types based on a Mann-Whitney U test (FDR-corrected P < 0.05).

We also found evidence that sex chromosome mismatch does not contribute to Y chromosome overexpression in late spermatogenesis in sterile *musculus*^♀^ × *domesticus*^♂^ hybrids. The Y chromosome was upregulated in *mus*×*dom* sterile hybrids relative to *dom*×*mus*^*dom*Y^ mice. This could be due to rescue of *domesticus* Y chromosome expression when paired with the *domesticus* X, but it could also be due to overall lower sex chromosome expression in mice with a *domesticus*^♀^ × *musculus*^♂^ background (Figure 5E). In Nonhybrid XY Mismatch, we saw that *mus*^*dom*Y^ mice had lower expression on the Y chromosome compared to *dom* controls, in contrast to the Y chromosome overexpression observed in *mus*×*dom* hybrids (Figure 5F). Thus, XY mismatch does appear to influence Y chromosome expression, but in the opposite direction of that observed in sterile hybrids.

In the reciprocal cross (*domesticus*^♀^ × *musculus*^♂^ F1 hybrids), we found some evidence that XY mismatch may contribute to disrupted expression of X-linked genes. Here Y chromosome expression was not different from that on the autosomes (Figure 5G), but the X chromosome tended to be downregulated (Figure 5J; Larson et al. 2017). There was no evidence that XY match restored normal X chromosome expression levels in *dom*×*mus*^*dom*Y^ (Hybrid F1 XY Match), with this cross type showing similar or even slightly lower expression levels on the X chromosome relative to *dom*×*mus* hybrids (Figure 5K). However, in Nonhybrid XY Mismatch we observed lower expression on the X chromosome in *dom*^*mus*Y^ mice relative to *dom* controls (Figure 5L). Therefore, a *domesticus* X paired with a *musculus* Y can result in suppression of X-linked gene expression even in the absence of autosomal incompatibilities.

### XY Mismatch Disrupted the Expression of Several Genes during Late Spermatogenesis

We also tested for effects of XY mismatch on individual genes by identifying differentially expressed (DE) genes in XY mismatch mice compared to controls. In our reanalysis, we identified many more overexpressed genes in sterile *mus*×*dom* hybrids compared to *mus* and many more underexpressed genes in the reciprocal *dom*×*mus* hybrids compared to *dom* on the X chromosome (Table 2), consistent with previous results (Larson et al. 2017) and with our observations of overall expression differences (Figure 5). We then asked if any of these X-linked DE genes were associated with XY mismatch. If so, then we would expect our Hybrid F1 XY Match *mus*×*dom*^*mus*Y^ to rescue some of the disrupted X-linked expression, and thus manifest as DE genes in comparisons between *mus*×*dom* and *mus*×*dom*^*mus*Y^. These genes should also overlap with genes DE between *mus*×*dom* and *mus*. However, there were only two X-linked DE genes in the *mus*×*dom* versus *mus*×*dom*^*mus*Y^ comparison (Table 2), and only one was also DE in the *mus*×*dom* versus *mus* comparison (Figure 6). This gene is a predicted protein coding gene, *Gm10058*, that shares 97% sequence identity with *Slx* and is therefore likely a paralog of this gene family. The other DE gene was *Btbd35f17*, another ampliconic gene with a protein-protein binding domain that is specifically expressed in male reproductive tissues (Smith et al. 2019). In Nonhybrid XY Mismatch, we only observed one X-linked DE gene in *mus*^*dom*Y^ compared to *mus*, and this gene was not DE in any other comparisons. Taken together, both Hybrid F1 XY Match and Nonhybrid XY Mismatch results suggest that almost all DE genes on the X chromosome in sterile *musculus*^♀^ × *domesticus*^♂^ hybrids are disrupted due to X-autosomal or autosomal-autosomal incompatibilities, rather than Y-linked incompatibilities.

**Table 2.** Number of differentially expressed genes in round spermatids for different cross types comparisons. “Higher” indicates higher expression (i.e., overexpressed) in the cross type with XY mismatch (F1 hybrids in Larson et al. 2017 and Hybrid F1 XY Match, Y-introgression F1 crosses in Nonhybrid XY Mismatch). “Lower” indicates lower expression (i.e., underexpressed) in the cross type with XY mismatch. For comparisons in the “Other Contrasts” category, “higher” indicates higher expression in the first cross type listed (*mus* or *mus×dom*). Gray boxes indicate chromosomes that are from the same subspecies in the two cross types being compared. Reciprocal F1s were considered as having the same autosomal backgrounds. Autosomal DE genes overlapping with putatively introgressed regions were excluded from comparisons involving Y-introgression mice.

|  |  | Autosomes |  | X Chromosome |  | Y Chromosome |  |
| --- | --- | --- | --- | --- | --- | --- | --- |
|  |  | Higher | Lower | Higher | Lower | Higher | Lower |
| Larson et al. 2017 | *mus*×*dom* vs *mus* | 1518 | 1476 | 252 | 13 | 109 | 66 |
|  | *mus*×*dom* vs *dom* | 1357 | 1241 | 190 | 55 | 15 | 2 |
|  | *dom*×*mus* vs *mus* | 1360 | 1009 | 62 | 73 | 6 | 8 |
|  | *dom*×*mus* vs *dom* | 1237 | 878 | 27 | 73 | 69 | 86 |
| Hybrid F1 XY Match | *mus*×*dom* vs *mus*×*dom*<sup>musY</sup> | 3 | 3 | 2 | 0 | 74 | 70 |
|  | *mus*×*dom* vs *dom*×*mus*<sup>domY</sup> | 21 | 83 | 38 | 96 | 68 | 84 |
|  | *dom*×*mus* vs *mus*×*dom*<sup>musY</sup> | 372 | 122 | 44 | 101 | 76 | 85 |
|  | *dom*×*mus* vs *dom*×*mus*<sup>domY</sup> | 2 | 4 | 1 | 0 | 71 | 66 |
| Non-hybrid XY Mismatch | *mus*<sup>domY</sup> vs *mus* | 13 | 34 | 1 | 0 | 52 | 63 |
|  | *mus*<sup>domY</sup> vs *dom* | 1820 | 2269 | 28 | 179 | 3 | 69 |
|  | *dom*<sup>musY</sup> vs *mus* | 1634 | 1679 | 70 | 55 | 10 | 7 |
|  | *dom*<sup>musY</sup> vs *dom* | 13 | 63 | 0 | 10 | 14 | 70 |
| Other Contrasts | *mus* vs *dom* | 1536 | 1774 | 38 | 139 | 46 | 96 |
|  | *mus*×*dom* vs *dom*×*mus* | 42 | 19 | 85 | 36 | 5 | 1 |

**Figure 6.**
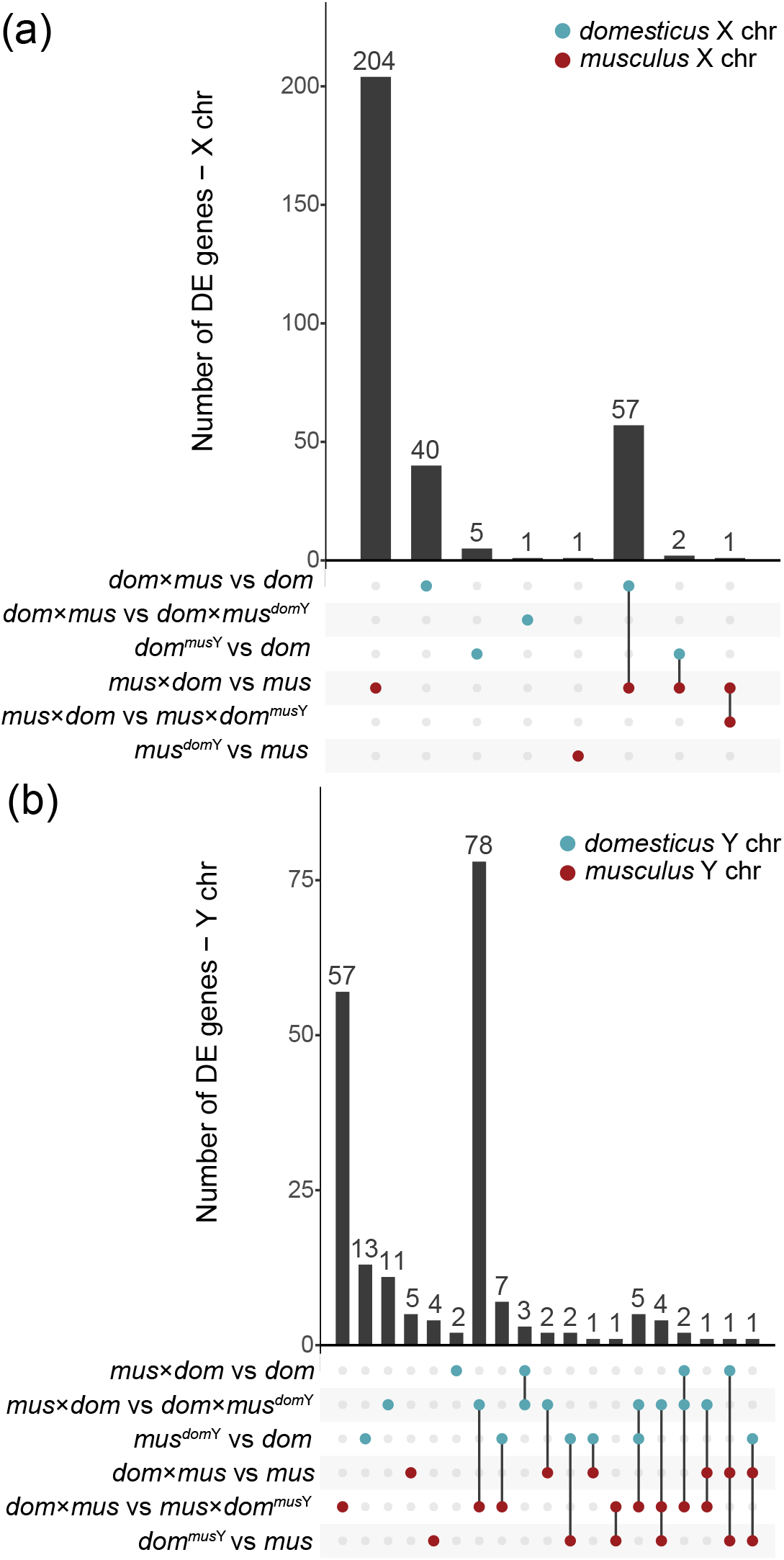
Upset plots showing the number of differentially expressed (DE) genes in each cross type comparison, and genes that are DE across multiple comparisons.(A) DE genes on the X chromosome. (B) DE genes on the Y chromosome. Bars corresponding to multiple dots connected by lines indicate genes that are DE across multiple comparisons. Bars corresponding to single dots indicate genes that are DE in only one comparison. The top three contrasts (blue dots) indicate comparisons on the *domesticus* X chromosome (A) or *domesticus* Y chromosome (B), and the bottom three contrasts (red dots) indicate comparisons on the *musculus* X chromosome (A) or *musculus* Y chromosome (B). Genes that were DE in opposite directions across multiple comparisons of the same sex chromosome were excluded.

On the X chromosome, very few DE genes were shared across multiple comparisons. However, 57 DE genes were shared between the *mus*×*dom* versus *mus* and *dom*×*mus* versus *dom* comparisons. When we looked at DE genes separated by direction of expression difference, only eight were shared between these two comparisons (Supplemental Material, Figure S7), so most of the overlap represented genes overexpressed in *mus*×*dom* but underexpressed in *dom*×*mus*. This could indicate that similar regulatory networks are disrupted in reciprocal F1 hybrids, but in ways that disrupt gene expression levels in opposite directions.

In contrast to the X chromosome, more Y-linked DE genes were shared across comparisons (Figure 6). Sterile *mus*×*dom* hybrids had 17 Y-linked DE genes that showed a clear bias towards overexpression (Table 2). Of these 17 DE genes, 5 were shared with the Hybrid F1 XY Match comparison *mus*×*dom* versus *dom*×*mus*^*dom*Y^, so having *domesticus* X and Y chromosomes partially rescued expression levels on the Y chromosome in *dom*×*mus*^*dom*Y^ mice. However, none of the 17 Y-linked genes DE in sterile hybrids were also DE in the Nonhybrid XY Mismatch comparison (*mus*^*dom*Y^ versus *dom*), so it is unlikely that XY mismatch alone disrupts expression of these genes. Instead, there may be a complex interaction between XY mismatch and a hybrid autosomal background that disrupts Y chromosome expression. Consistent with this, we found the most Y-linked DE genes in comparisons between cross types with reciprocal hybrid autosomal backgrounds but the same Y chromosome (Table 2). Of these, 78 Y-linked DE genes were shared between these two comparisons (Figure 6), suggesting that reciprocal hybrid autosomal backgrounds may have resulted in disrupted expression for many of the same Y-linked genes, regardless of the subspecies origin of the Y chromosome.

We also found several autosomal genes that were DE between cross types with the same autosomal background but different sex chromosome combinations (Table 2). We excluded autosomal genes that overlapped with putatively introgressed regions, so the DE that we detected was unlikely to result from *cis*-regulatory effects of variants from the opposite subspecies that introgressed along with the Y chromosome. In Hybrid F1 XY Match, 104 autosomal genes were DE when comparing *mus*×*dom* to *dom*×*mus*^*dom*Y^ and 494 autosomal genes were DE when comparing *dom*×*mus* to *mus*×*dom*^*mus*Y^ (Table 2). These comparisons involved reciprocal crosses with the same autosomal and Y chromosome genotypes, and so DE presumably resulted from X-autosomal incompatibilities. Although overexpression on the X chromosome tends to be the most notable expression pattern associated with X-autosomal incompatibilities, previous studies have shown disrupted postmeiotic autosomal expression in sterile hybrids as well (Larson et al. 2017). We detected only six (non-overlapping) DE genes in each comparison with different Y chromosomes but the same autosomal and X chromosome genotypes (*mus*×*dom* versus *mus*×*dom*^*mus*Y^ and *dom*×*mus* versus *dom*×*mus*^*dom*Y^; Table 2).

In Nonhybrid XY Mismatch, we identified some autosomal DE genes in comparisons that had different Y chromosomes but the same autosomal and X backgrounds, suggesting that interactions involving the Y chromosome disrupted some autosomal expression, but the number of autosomal DE genes was not enriched relative to the number of X-linked DE genes (Fisher’s Exact Test P > 0.05; Table 2). These autosomal DE genes tended to be underexpressed in the cross type with XY mismatch regardless of the direction of the cross (Table 2) and must result from direct interactions with the Y chromosome or indirect interactions with XY mediated expression changes. Only one autosomal gene, *Babam2*, was DE in both reciprocal comparisons. It is a member of the BRCA1-A complex, which is involved in DNA double-strand break repair (The Uniprot Consortium 2020).

Finally, we tested if DE genes tended to be in the same co-expression networks using weighted correlation network analysis (WGCNA). We found one module in Hybrid F1 XY Match associated with the *mus*×*dom* autosomal background, one module in Nonhybrid XY Mismatch associated with the *musculus* background, and one module in Nonhybrid XY Mismatch associated with the *domesticus* background (Figure 7A, B, D). These modules were significantly enriched for genes DE between cross types with different autosomal backgrounds (Table 3). There were also multiple modules enriched for DE genes despite not having a significant association with cross type (Table 3). For example, Module 5 was significantly enriched for DE genes in all pairwise comparisons in Hybrid F1 XY Match. Although we did not detect a significant cross type association for this module, there was a trend towards an autosomal background by sex chromosome effect for this module, with *mus*×*dom* background cross types tending to have lower module membership in general, but with *mus*×*dom*^*mus*Y^ mice tending to have higher module membership than *mus*×*dom* mice (Figure 7E). Another Hybrid F1 XY Match module showed a similar pattern (Module 3, Figure 7C) and was enriched for genes DE between *dom*×*mus* and *mus*×*dom*^*mus*Y^ (Table 3). In Nonhybrid XY Mismatch, Module 5 was enriched for genes DE between *mus*^*dom*Y^ and either subspecies (*mus* or *dom*; Table 3), and XY mismatch mice tended to have lower associations with this module (Figure 7). We likely did not have enough power to detect significant module associations with complex autosome by sex chromosome interactions given our sample size, especially because these effects on gene expression tended to be subtle and affect relatively few genes (Figure 5, Table 2). Despite low power, the fact that certain modules were enriched for DE genes suggests that groups of genes were disrupted in similar ways in XY mismatch mice, and that particular gene networks may be disrupted under XY mismatch. Additionally, we found a significant positive correlation in module eigengene values between Hybrid F1 XY Match and Nonhybrid XY Mismatch (Module 5 in both experiments, r = 0.64; FDR-corrected Pearson’s correlation P < 0.001; Supplemental Figure S8) and a significant overlap in genes (279 genes, FDR-corrected Fisher’s Exact Test P < 0.001), suggesting that these two modules represent genes with similar expression patterns between the two experiments. Interestingly, these modules trended towards a negative association with cross types that had a *musculus* X chromosome and *domesticus* Y chromosome (Figure 7E, 7F), and may represent genes with similar expression patterns under XY mismatch regardless of autosomal background. All DE genes and their module memberships are listed in Supplemental Material, Tables S6 and S7.

**Figure 7.**
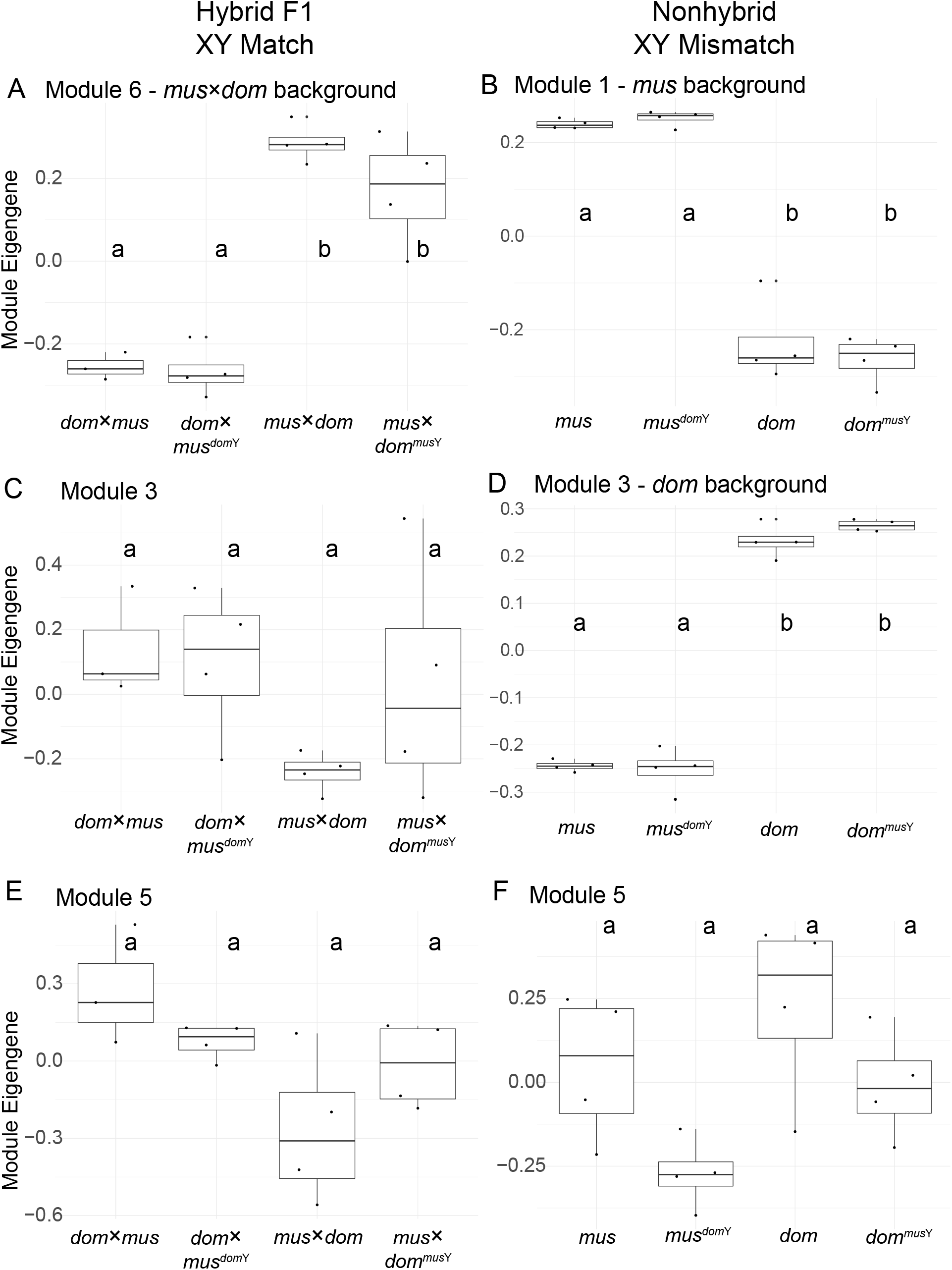
Example WGCNA module eigengene values plotted by cross type. Note that WGCNA was performed separately for each experiment, so there is not necessarily a relationship between Hybrid F1 XY Match and Nonhybrid XY Mismatch modules with the same number. Modules that were significantly associated with cross types are also labeled based on these associations (A, B, and D). Other modules shown were not significantly associated with a cross type but trended towards an association with X-autosomal background by Y chromosome type interaction and were enriched for DE genes in at least one comparison (C, E, and F; Table 3). Letters indicate significant differences in module association based on linear models with post-hoc Tukey tests (P < 0.05).

**Table 3:**
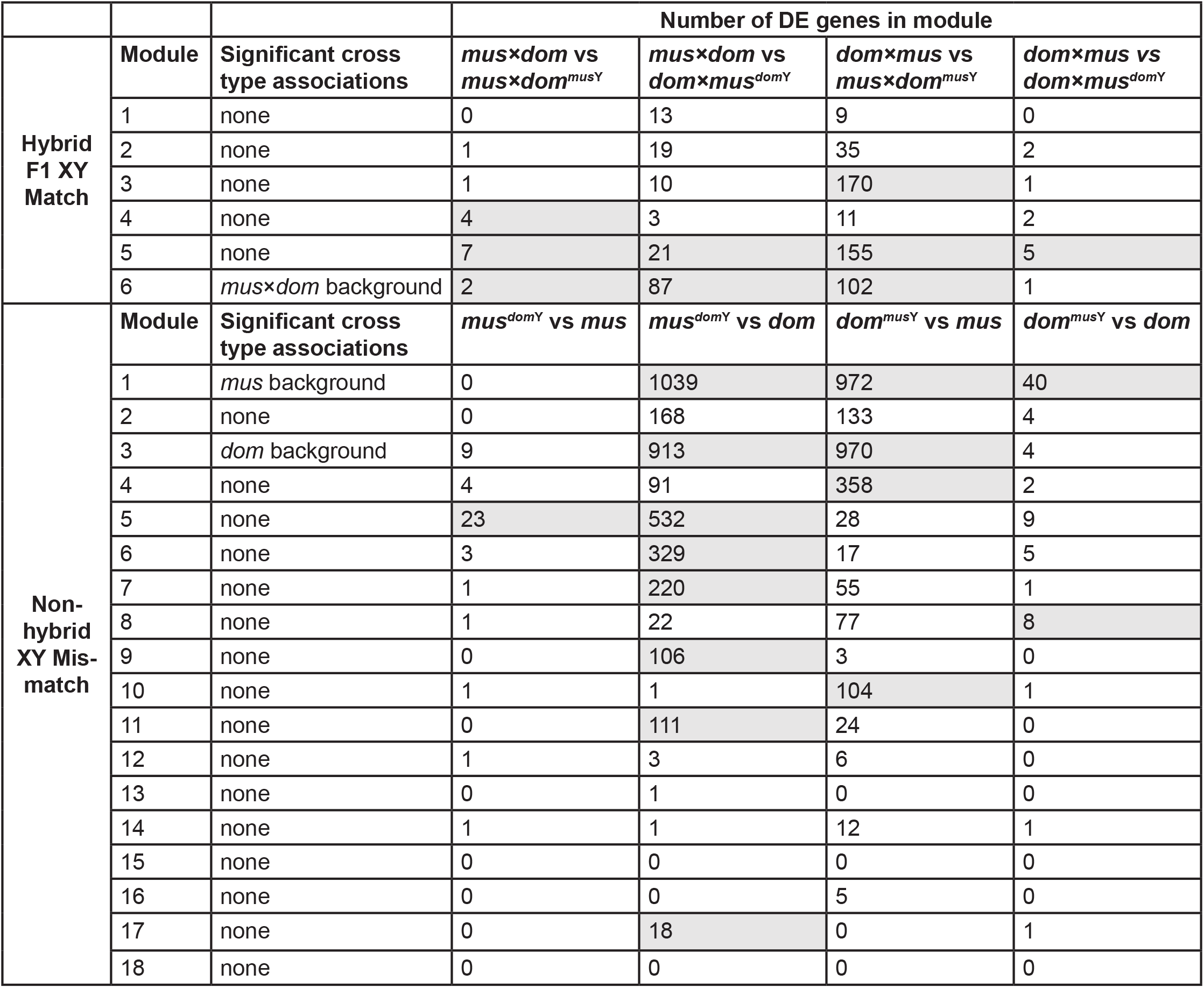
Number of differentially expressed genes in each WGCNA module. Rows indicate WGCNA modules and columns indicate comparisons between cross types used to identify differentially expressed (DE) genes. Module associations with cross types are based on linear models with post-hoc Tukey tests. Shaded boxes indicate a significant enrichment for DE genes based on a hypergeometric test with FDR-correction (P < 0.05). Note that there is not necessarily a relationship between Hybrid F1 XY Match and Nonhybrid XY Mismatch modules with the same module number.

## Discussion

The large X-effect and Haldane’s rule are prevalent patterns observed in intrinsic hybrid incompatibilities across diverse taxa and suggest that sex chromosomes play a predominant role in speciation, but the evolutionary forces underlying rapid sex chromosome divergence that leads to hybrid incompatibilities remain unclear (Presgraves and Meiklejohn 2021). One compelling hypothesis is that hybrid incompatibilities are a consequence of intragenomic conflict between sex chromosomes (Frank 1991; Hurst and Pomiankowski 1991; Lindholm et al. 2016). In this study, we showed that intragenomic conflict between the sex chromosomes may contribute to some hybrid incompatibilities in house mice, but not in a simple dosage-dependent manner, and with subtle effects relative to other components of F1 hybrid incompatibilities. Notably, we find that XY conflict does not appear to contribute to postmeiotic disruption of sex chromosome repression, a major regulatory phenotype associated with hybrid sterility in house mice (Larson et al. 2017). Below, we discuss the implications of our findings for the genetic basis of house mouse male hybrid sterility and the potential role of intragenomic conflict in speciation.

### Insights into the Genetic Basis of Mouse Male Hybrid Sterility

Our results did not support the model of *Slx*- and *Slxl1*-*Sly* dosage imbalance leading to X chromosome overexpression in mouse F1 hybrids. In Hybrid F1 XY Match, we showed that XY match on an F1 background did not restore postmeiotic X chromosome repression (Figure 5). In Non-hybrid XY Mismatch, we directly tested the effects of XY mismatch in the absence of X-autosomal mismatch on post-meiotic spermatogenesis gene expression. We found some evidence for disrupted expression in XY mismatch mice (Figure 5, Table 2), but the effects were relatively subtle and often in the opposite direction than expected based on genetic manipulation studies (Cocquet et al. 2012; Kruger et al. 2019) or disrupted expression in sterile F1 mice (Larson et al. 2017; Figures 4, 5, and 6).

Our results indicate that genetic manipulation studies, which performed nearly complete knockdowns or duplications, are not representative of the more subtle copy number differences expected to occur in natural hybrids (Cocquet et al. 2009; Cocquet et al. 2012; Kruger et al. 2019). Another important difference from genetic manipulation studies is that we used wild-derived inbred strains instead of the C57BL/6J classic laboratory mouse, which has a mostly *domesticus* background but some *musculus* introgression throughout, including the Y chromosome (Nagamine et al. 1992). Because C57BL/6J is mostly *domesticus* with a *musculus* Y chromosome, it has a similar genetic composition as our wild-derived *dom*^*mus*Y^ mice and therefore may show some of the same subtle disruptions to gene expression and sperm morphology that we observed compared to pure *domesticus* mice. We also introgressed the entire Y chromosome, so there should not have been dosage imbalances among ampliconic genes on the same sex chromosome. However, our Y-introgression mice also had imbalance between all Y-linked ampliconic genes and interacting genes on the X chromosome and autosomes, so it is unclear if introgressing the entire Y chromosome should cause larger or smaller effects on postmeiotic spermatogenesis expression.

SLX, SLXL1, and SLY proteins interact with other sex-linked and autosomal ampliconic genes, including *Ssty1*/2, *α*-*takusan*, and *Speer*, so additional gene families may be involved in intragenomic conflict with *Slx, Slxl1*, and *Sly* (Kruger et al. 2019; Moretti et al. 2020). Our autosomal gene family expression results seem to further complicate understanding of the consequences of ampliconic gene conflict as we found that the *α*-*takusan* gene family is overexpressed in F1 hybrids regardless of cross direction or sex chromosome type (Figure 4F). Sex chromosome mismatch, however, did not disrupt *α*-*takusan* expression when the autosomal background was nonhybrid. This was somewhat puzzling because protein products of sex-linked ampliconic genes are thought to regulate *α*-*takusan* expression in late spermatogenesis, perhaps again indicating that copy number differences between subspecies are too subtle to generate strong regulatory phenotypes. Another surprising expression result was that *Slxl1* expression levels were not correlated with *Slxl1* copy numbers (Figure 4B). Other genes are likely involved in the regulation of *Slxl1* (Moretti et al. 2020), and it is possible that the evolution of these *trans*-acting factors may play a more important role in determining overall *Slxl1* expression levels than *Slxl1* copy number *per se*.

On balance, our results suggest that differences in *Slx*- or *Slxl1*-*Sly* dosage do not result in strong hybrid incompatibilities. We did not observe sex chromosome overexpression with an excess of *Slx* and *Slxl1* copies or underexpression with an excess of *Sly* copies as predicted under the conflict model (Larson et al. 2017). Therefore, the primary mechanisms underlying postmeiotic X chromosome overexpression in sterile F1 hybrids likely do not involve XY interactions. Instead, disrupted postmeiotic repression is likely a continuation of *Prdm9*-mediated MSCI disruption (Bhattacharyya et al. 2013; Bhattacharyya et al. 2014; Mukaj et al. 2020).

Although XY copy number imbalance is unlikely to explain disrupted postmeiotic repression in F1 hybrids, sex chromosome interactions may play a role in house mouse hybrid sterility. We showed that XY mismatch can lead to disrupted expression of ampliconic genes and other genes throughout the genome (Figure 4, Figure 6, Table 2), and some of these genes are essential for spermatogenesis. For example, *Taf7l* knockouts have abnormal sperm morphology (Cheng et al. 2007), *Prdx4* knockouts have reduced sperm counts (Iuchi et al. 2009), and both these genes were differentially expressed in *dom*^*mus*Y^ mice. We also showed that hybrid interactions involving the Y-chromosome are associated with subfertility phenotypes (Table 1), consistent with previous studies (Campbell et al. 2012; Campbell and Nachman 2014). Here we have focused on interactions between the sex chromosomes because the ampliconic gene conflict model established a clear prediction for XY incompatibilities, but we could not distinguish XY incompatibilities from Y-autosomal incompatibilities in our experimental crosses. We note that several of our observations could result from Y-autosomal interactions. Indeed, introgression of the Y chromosome (Nonhybrid XY Mismatch) induced autosomal regulatory phenotypes.

We observed some autosomal regions that co-introgressed with the Y chromosome, and some of these regions have been shown to introgress in other mouse hybrids (Supplemental Material, Figure S4). These may be regions that are incompatible with the Y chromosome from the opposite subspecies, and therefore must co-introgress for mice to be viable or fertile. The large introgressed region we identified on chromosome 2 is adjacent to a multicopy gene, *R2d2*, involved in meiotic drive during female meiosis (Didion et al. 2016). *R2d2* has only been shown to act in females (Didion et al. 2016), but our crossing scheme only involved backcrossing hybrid males. We also generated Y-intogression mice using the LEWES/EiJ strain, which is fixed for the low copy number allele of *R2d2*, and PWK/PhJ, which also appears to have low *R2d2* copy number (Didion et al. 2016), so it is unlikely that this introgression is a direct result of *R2d2* drive as previously described. Nevertheless, the exact functions of *R2d2* are unresolved, so this large region of introgression may be related to *R2d2*, but probably not through a direct meiotic drive mechanism.

Our results are likely important in the context of mouse speciation in nature. Mice sampled from the European hybrid zone are often advanced generation hybrids with complex patterns of ancestry from both *musculus* and *domesticus*, and true F1 genotypes are exceptionally rare (Teeter et al. 2010; Turner et al. 2012). Therefore, understanding mechanisms of hybrid incompatibility in addition to F1 X-autosomal incompatibilities is essential for understanding the complex genetic basis of mouse speciation occurring in nature. The Nonhybrid XY Mismatch experiment demonstrated that disrupted gene expression phenotypes can occur in the absence of an F1 autosomal background. Previous studies have shown that advanced intercrosses of hybrid mice show different sterility phenotypes than F1s (Campbell et al. 2012), and *Prdm9*-mediated hybrid sterility requires an F1 autosomal background, leading others to speculate that genetic incompatibilities underlying hybrid sterility may be different in later hybrid generations (Campbell and Nachman 2014; Mukaj et al. 2020). Our results show that Y chromosome introgression can contribute to reduced fertility (consistent with Campbell et al. 2012) and some disrupted spermatogenesis gene expression in later generation hybrids with non-F1 autosomal backgrounds.

### What Is the Contribution of Sex Chromosome Conflict to Speciation?

Several studies have proposed a link between intragenomic conflict and hybrid incompatibilities (Tao et al. 2001; Phadnis and Orr 2009; Wilkinson et al. 2014; Zanders et al. 2014; Case et al. 2015; Zhang et al. 2015; Larson et al. 2017), but it remains unknown how prevalent these systems are in natural populations or if intragenomic conflict is the primary selective force behind the evolution of these incompatibilities. While X-autosomal incompatibilities are known to play a central role in house mouse hybrid sterility, previous work has shown that house mouse speciation likely has a more complex genetic basis (Vyskočilová et al. 2005; Good et al. 2008b; Turner et al. 2012; Turner and Harr 2014; Larson et al. 2018b) and may involve sex chromosome intragenomic conflict (Ellis et al. 2011; Campbell et al. 2012; Larson et al. 2017). The exact mechanisms underlying reduced fertility associated with Y chromosome mismatch is unknown, and it is still unclear what role, if any, sex chromosome intragenomic conflict may play (Ellis et al. 2011; Campbell et al. 2012; Larson et al. 2017).

Ampliconic genes are a common feature of mammalian sex chromosomes, and they tend to be expressed specifically during spermatogenesis (Li et al. 2013; Soh et al. 2014; Skinner et al. 2016; Lucotte et al. 2018; Bellott et al. 2017; Hughes et al. 2020; reviewed in Larson et al. 2018a). Although difficult to quantify, evolution of ampliconic gene families involved in spermatogenesis is arguably one of the most rapidly evolving components of mammalian genomes (Mueller et al. 2013; Soh et al. 2014; Lucotte et al. 2018; Cechova et al. 2020; Vegesna et al. 2020). Intragenomic conflict among sex chromosome ampliconic genes has been proposed as a mechanism through which hybrid incompatibilities have evolved in at least three mammalian groups (cats, Davis et al. 2015; primates, Dutheil et al. 2015; and mice, Larson et al. 2018a; Kruger et al. 2019). In cats, loci associated with hybrid sterility tend to be in or near high copy number genes (Davis et al. 2015). In great apes, sex chromosome amplicon copy number can evolve rapidly (Lucotte et al. 2018; Cechova et al. 2020), and ampliconic regions on the X chromosome are thought to have experienced selective sweeps as a result of strong selection pressures imposed by intragenomic conflict with the Y chromosome (Nam et al. 2015). These regions also overlap sections of the modern human X chromosome that lack Neandertal introgression, and therefore may represent regions involved in genetic incompatibilities between modern humans and Neandertals (Dutheil et al. 2015). However, most of these connections remain speculative and the X chromosome is clearly a hotspot of the evolution of hybrid incompatibilities (Masly and Presgraves 2007; Good et al. 2008a).

Theoretical work introducing the idea that sex chromosome intragenomic conflict could contribute to hybrid incompatibilities focused on this phenomenon as an explanation for Haldane’s rule and the large X-effect (Frank 1991; Hurst and Pomiankowski 1991). However, genetic conflict between the sex chromosomes during reproduction cannot explain some observations, such as the applicability of Haldane’s rule and the large X-effect to hybrid inviability or the important role of the X chromosome in many incompatibilities that occur in homogametic hybrids (Coyne 1992). In this study, we showed that XY conflict may have a small effect on male hybrid sterility, but *Prdm9*-mediated incompatibilities probably play the most important role in the observations consistent with Haldane’s rule and the large X-effect in house mice. Interactions among *Prdm9, Hstx2*, and other autosomal and X-linked loci in hybrids result in failed or delayed double strand break repair, which eventually leads to meiotic arrest and male sterility (Forejt et al. 2021). The rapid divergence of *Prdm9* and its binding sites is likely the result of PRDM9 haplotype selection, leading to biased gene conversion and hotspot erosion (Baker et al. 2015). Thus, intragenomic conflict is unlikely to be the primary underlying cause of house mouse hybrid male sterility.

It remains unknown if the recurrent evolution of ampliconic genes is a consequence of intragenomic conflict across mammals, but this is generally assumed to be the case. If so, intragenomic conflict may be much more important in the evolution of hybrid incompatibility loci than once thought (Johnson and Wu 1992; Coyne and Orr 2004). Some recent empirical studies support this hypothesis in both flies and mammals (Presgraves and Meiklejohn 2021), however, our study did not provide direct support for this hypothesis. XY mismatch likely contributes to hybrid male sterility and disrupted expression, but in more complex ways than the *Slx, Slxl1*, and *Sly* dosage-based conflict model, and with relatively small effects on hybrid sterility. In particular, we note that *Ssty1* and *Sly* show opposing copy number patterns between subspecies, such that replacing a *musculus* Y with a *domesticus* Y simultaneously increases *Ssty1* while decreasing *Sly*, and vice versa. It is possible that higher copy number of one gene can compensate for reduced copy number of the other in regulating postmeiotic sex chromatin. This work is thus distinct from previous work focused on deletions (that reduce copy number of both genes) or RNA interference (that selectively targets one gene).

Further work is required to identify loci involved in XY or Y-autosomal incompatibilities, but it is plausible that intragenomic conflict among ampliconic genes still plays a role given that these genes are the primary sex chromosome genes expressed in the postmeiotic stages during which spermatogenesis expression is highly disrupted (Sin and Namekawa 2013; Larson et al. 2017). Copy number mismatch between these gene families may play important roles in reproductive outcomes in nature, as has been implied from slight sex ratio skews in regions of the hybrid zone (Macholán et al. 2008). Even subtle differences in fertility could have important effects on fitness, especially given that sperm competition appears to be common in mice (Dean et al. 2006). However, our work suggests that such effects do not manifest as a major reproductive barrier between populations.

## Supporting information

KopaniaEtal_supplement

## Data Availability

Whole genome sequence data from Y-introgression strains and RNAseq data from testes cell sort populations are publicly available through the National Center for Biotechnology Information Sequence Read Archive under accession numbers PRJNA816542 (whole genome) and PRJNA816886 (RNAseq). Raw phenotype data are available in the Supplemental Material, Table S2. Scripts used to modify the AmpliCoNE program for copy number estimation are publicly available at: https://github.com/ekopania/modified-AmpliCoNE. Scripts used for gene expression analyses are available at: https://github.com/ekopania/xy_mismatch_expression_analyses.

## Acknowledgements

We would like to thank Sara Keeble for assistance with animal husbandry, Pamela K. Shaw and the UM Fluorescence Cytometry Core supported by an Institutional Development Award from the NIGMS (P30GM103338 and S10-OD025019) and the UM Lab Animal Resources staff. This study included research conducted in the University of Montana Genomics Core, supported by a grant from the M. J. Murdock Charitable Trust (to J.M.G.). Computational resources and support from the University of Montana’s Griz Shared Computing Cluster (GSCC), supported by grants from the National Science Foundation (CC-2018112 and OAC-1925267, J.M.G. co-PI), contributed to this research. We thank members of the Good Lab, Polly Campbell, Lila Fishman, Doug Emlen, and Travis Wheeler for helpful discussions and comments during the development of this study. We thank three anonymous reviewers for helpful comments on an earlier version of this manuscript. This work was supported by grants from the Eunice Kennedy Shriver National Institute of Child Health and Human Development of the National Institutes of Health (R01-HD073439, R01-HD094787 to J.M.G.). E.E.K.K. was supported by the National Science Foundation Graduate Research Fellowship Program (DGE-1313190), a grant from the National Science Foundation (DEB-1754096 to J.M.G.), and a Rosemary Grant Award for Graduate Student Research from the Society for the Study of Evolution. E.L.L. was supported by the National Science Foundation (DEB-2012041). C.C.R. was supported by the BBSRC (BB/N000463/1 to P.E.). P.E. also acknowledges funding from the Leverhulme Trust (RPG-2019-414 194). E.M.W. was supported by a faculty studentship from the University of Essex. B.M.S was supported by UKRI (University of Essex). Any opinions, findings, and conclusions or recommendations expressed in this material are those of the author(s) and do not necessarily reflect the views of the National Science Foundation, the National Institutes of Health, or the Society for the Study of Evolution.

## Author Contributions

J.M.G. and E.L.L. conceived the project. E.E.K.K., E.L.L., and J.M.G. designed the experiments. E.L.L. and E.E.K.K. did the mouse husbandry and breeding. E.E.K.K. performed the mouse dissections, cell sorts, and sequencing library preparation. E.M.W., C.C.R., B.M.S., and P.J.I.E. performed and analyzed the sperm morphology assays. E.E.K.K. analyzed the data. E.E.K.K., E.L.L., and J.M.G. wrote the manuscript with input from all authors.

### Conflicts of Interest

The authors declare no conflicts of interest.

## Notes

### Competing Interest Statement

The authors have declared no competing interest.

### Summary of Updates

Additional background information and discussion; Figures revised for clarity

