## Supplementary material for "The contribution of sex chromosome conflict to disrupted spermatogenesis in hybrid house mice": KopaniaEtal_supplement

**Supplementary Figures**

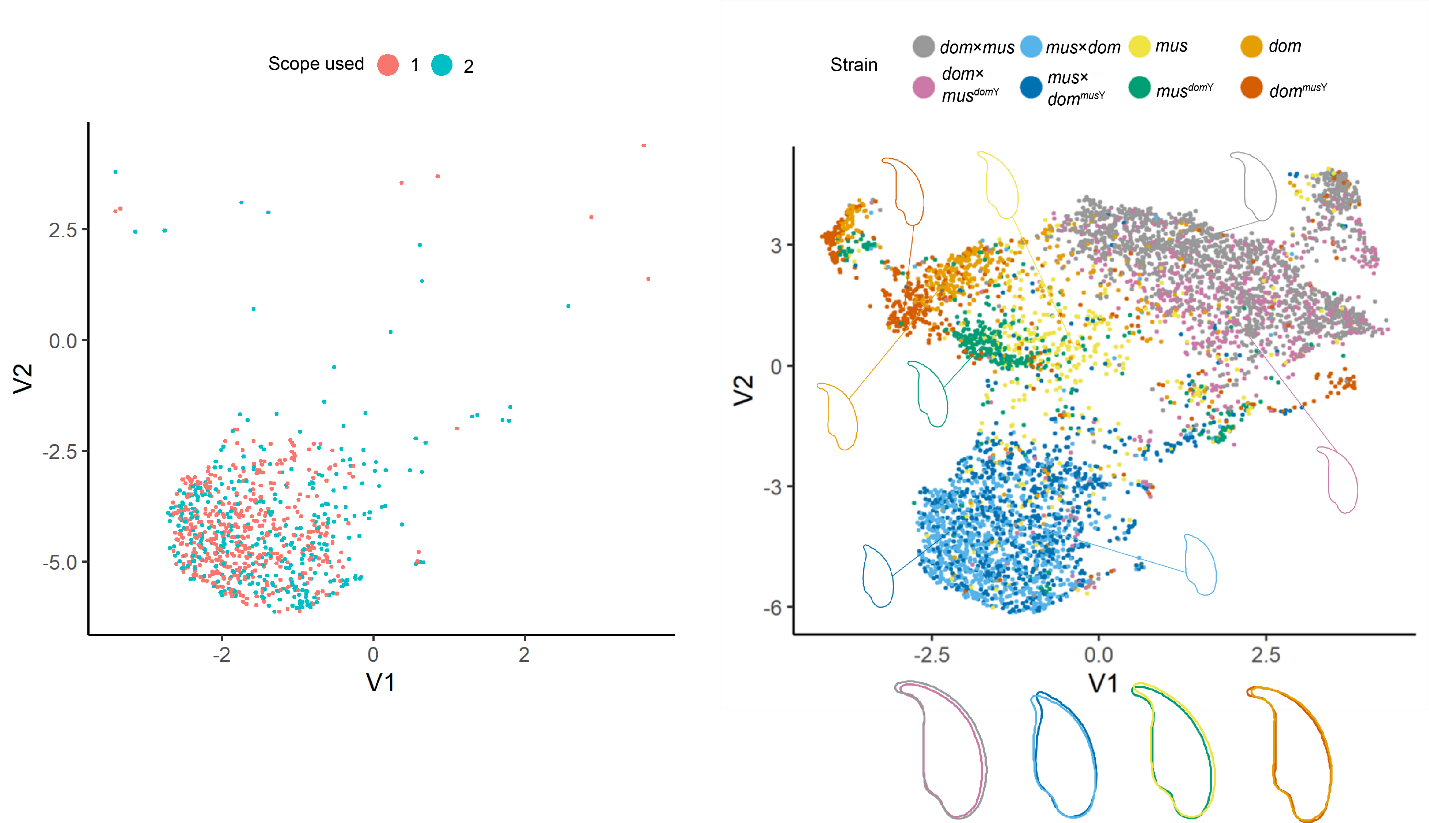

**Figure S1:** Dimensionality reduction plots (UMAP) of sperm nuclei morphology. (A) Nuclei from the *mus*×*dom* samples, which were imaged using two different microscopes, are evenly distributed within their cluster regardless of which microscope was used. This shows us that there was no experimental bias occurring based on which microscope was used, which allowed consistent data collection of the remaining samples from both scopes. (B) Clustering for all imaged nuclei colored by cross type.

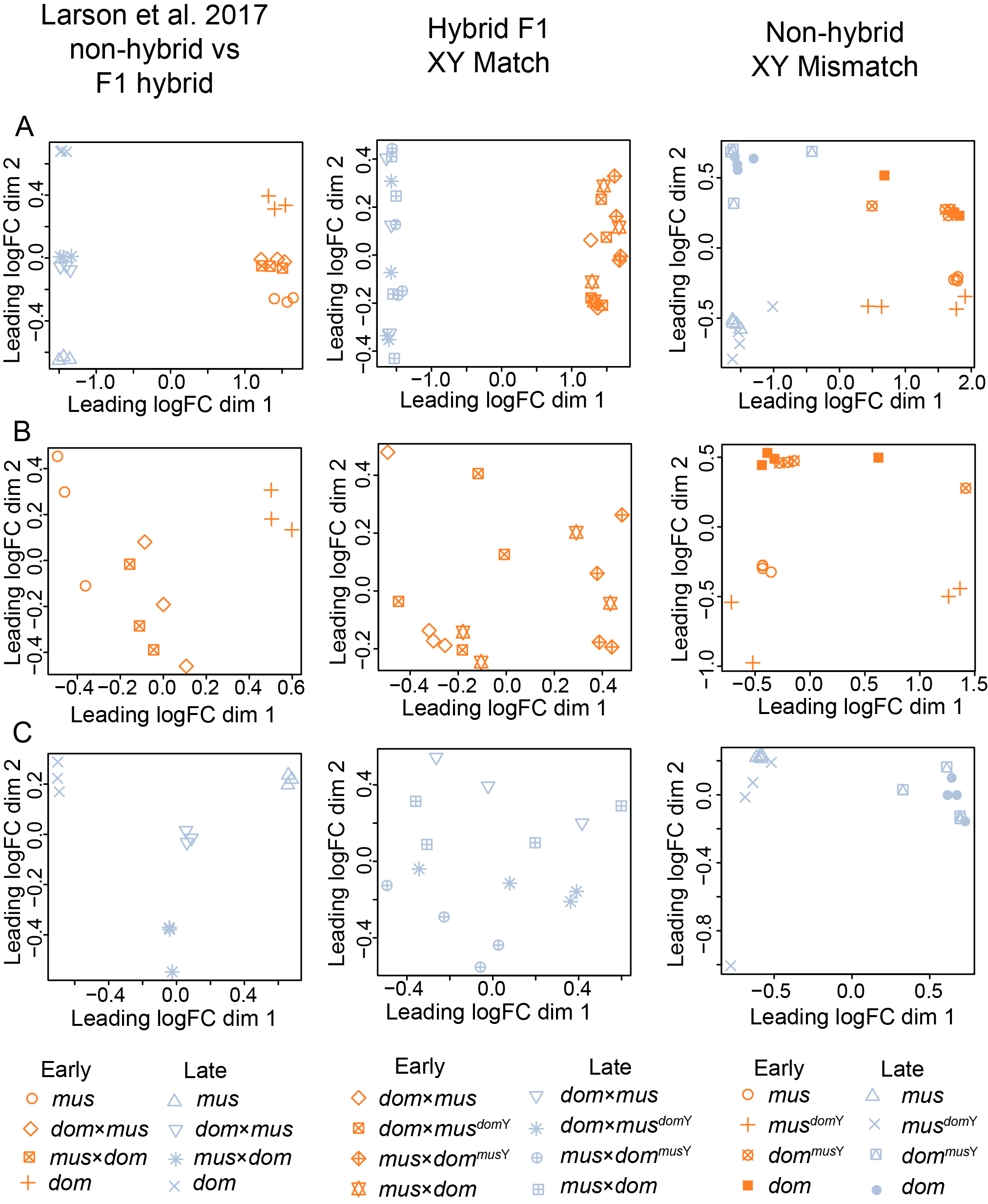

**Figure S2:** Multidimensional scaling (MDS) plots of distances between expression data from sorted cells. (A) Data from leptotene-zygotene (LZ, orange) and round spermatids (RS, blue) combined. (B) Data from LZ only. (C) Data from RS only. The first column shows data from our reanalysis of data from (Larson, et al. 2017). The second two columns show data collected from our Hybrid F1 XY Match and Non-hybrid XY Mismatch experiments. Each shape represents a different cross type, which are different across experiments. See legend at the bottom of the figure.

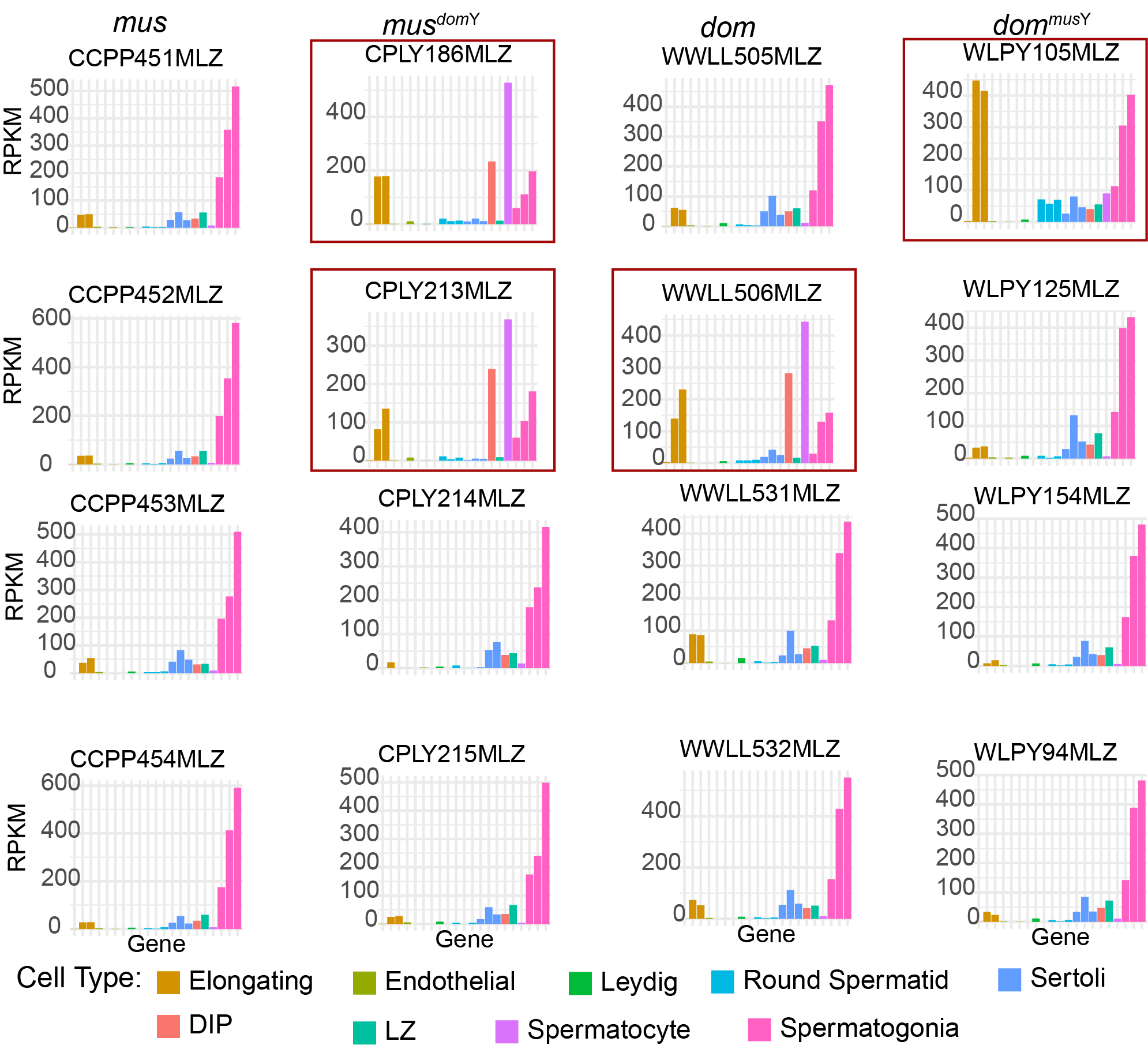

**Figure S3:** Expression levels of cell type marker genes in Non-hybrid XY Mismatch leptotene-zygotene (LZ) samples. Each plot represents a different individual, labeled by sample ID. Each column contains samples from the same cross type. Marker genes on the x-axis are colored by the cell type they are preferentially expressed in, and are based on single-cell RNAseq data (Green, et al. 2018). The y-axis indicates gene expression level in FPKM. LZ are known to have similar expression profiles to spermatogonia, so the high expression levels of spermatogonia marker genes is expected (Larson, et al. 2016). Note that the absolute expression level of these marker genes in the cell types they represent is highly variable, and a previous study showed that the LZ marker gene has a median FPKM value of about 25, so the relatively low FPKM value for the LZ marker in these plots is also expected (Hunnicutt, et al. 2021). Red boxes indicate samples that appear to have contamination from diplotene and elongating spermatid cells based on their relative expression levels of marker genes for the cell types compared to other LZ samples. These are the same 4 samples that separate from other LZ samples on leading logFC dim in Figure S1A and B (3^rd^ column), indicating that enough cell type contamination occurred in these samples to affect their overall expression profiles.

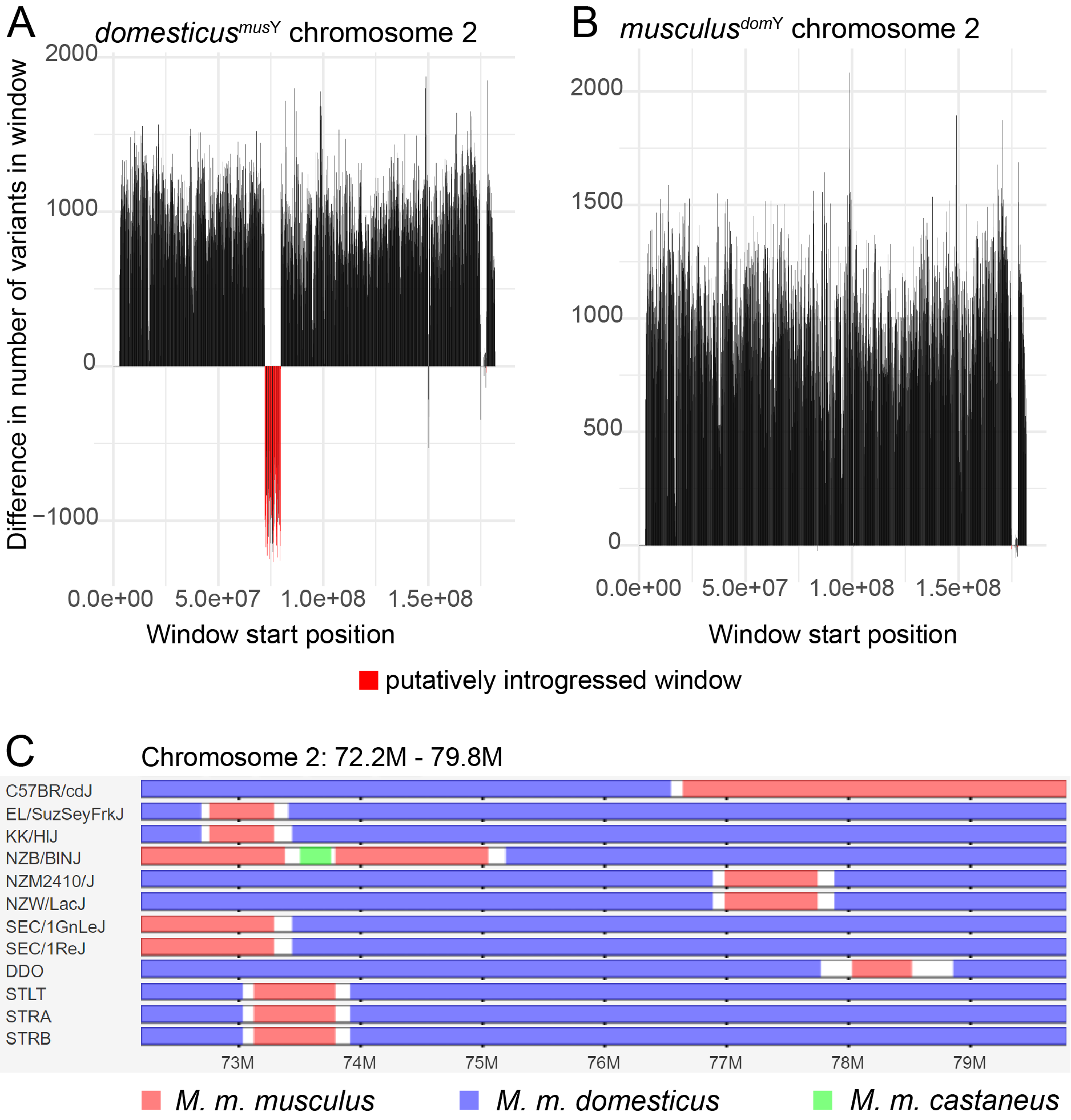

**Figure S4:** Evidence for introgression from *musculus* into *domesticus* on chromosome 2. (A) and (B) show the difference in the number of variants when Y introgression strains were mapped to their Y chromosome origin reference genome compared to their autosomal background reference genome in 100kb windows. Regions with evidence for introgression had more variants compared to the autosomal background reference than compared to the Y chromosome reference and are shown in red. Chromosome 2 has a large introgressed region in *domesticus^mus^*^Y^ (A) but not *musculus^dom^*^Y^ (B). (C) shows a screenshot from the Mouse Phylogeny Viewer (Yang, et al. 2011) depicting mouse inbred strains with evidence for introgression from *musculus* (red) into *domesticus* (blue) in the region of chromosome 2 where we found evidence for introgression.

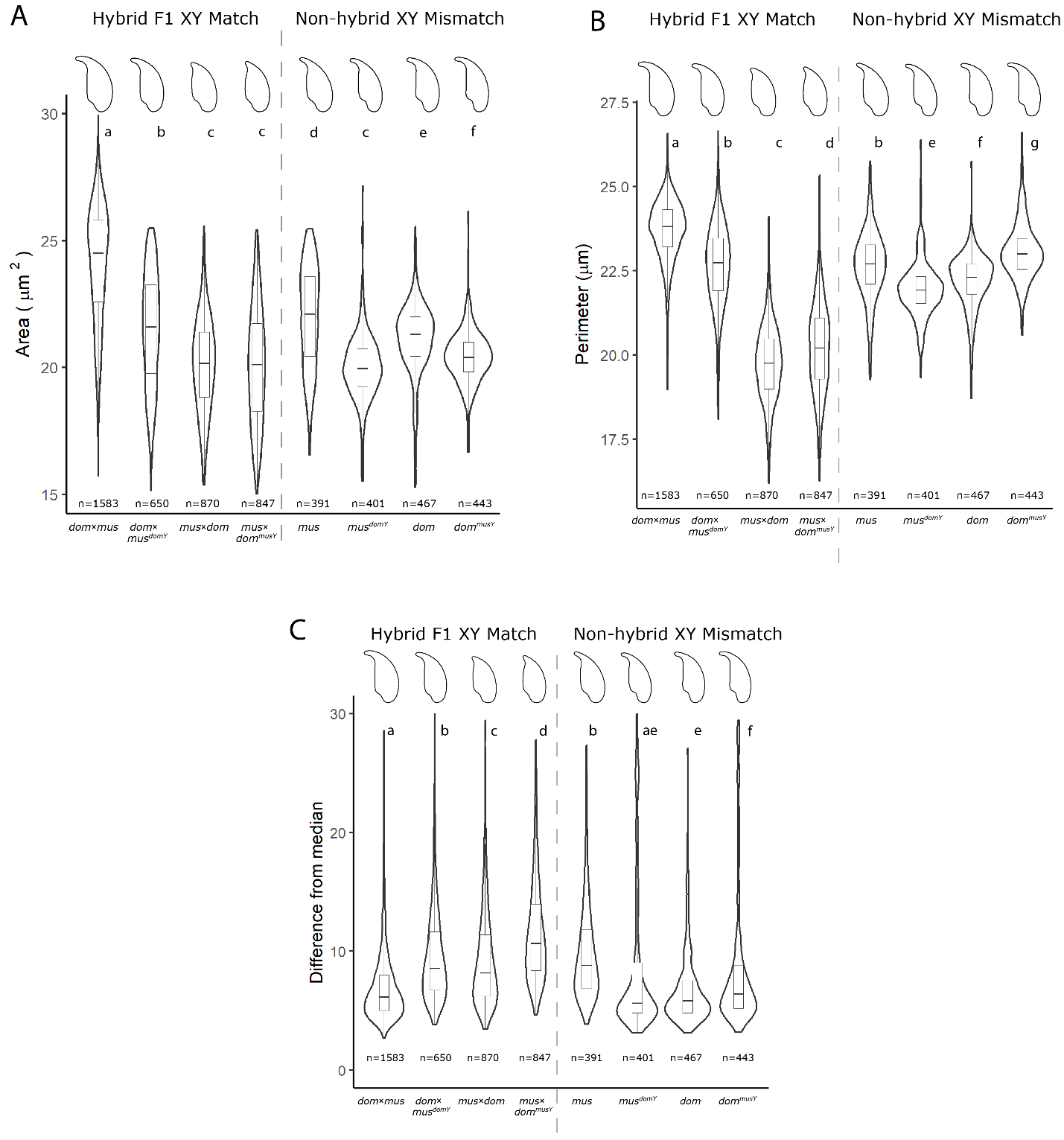

**Figure S5:** Violin plots showing sperm nuclear morphology parameters: (A) area (µm^2^), (B) perimeter (µm), and (C) difference from median. Difference from median is a measure of variance within cross types. Letters above each violin plot indicate significant differences among cross types based on an FDR-corrected pairwise Wilcoxon rank sum test. Numbers below each violin plot represent the number of sperm head nuclei observed for each cross type.

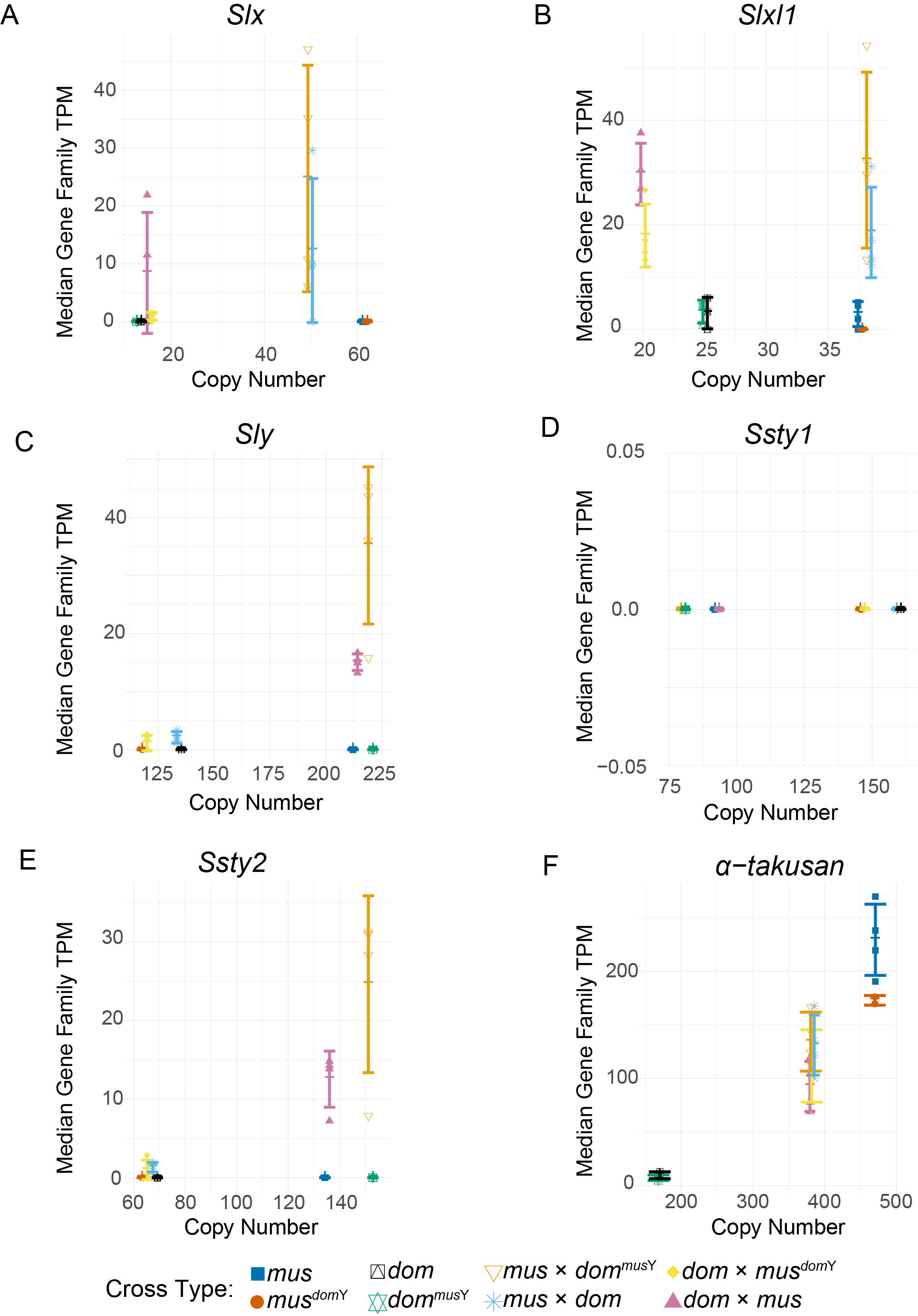

**Figure S6:** Normalized expression levels in leptotene-zygotene of *Slx* (A), *Slxl1* (B), *Sly* (C), *Ssty1* (D), *Ssty2* (E), and *𝛼-takusan* (F) ampliconic gene families in different cross types plotted against their copy numbers. Expression level was calculated by summing transcripts-per million (TPM) for each paralog of the gene family with at least 97% sequence identity to the ampliconic gene. Points represent values for individual samples, and lines indicate median and standard deviation for each cross type.

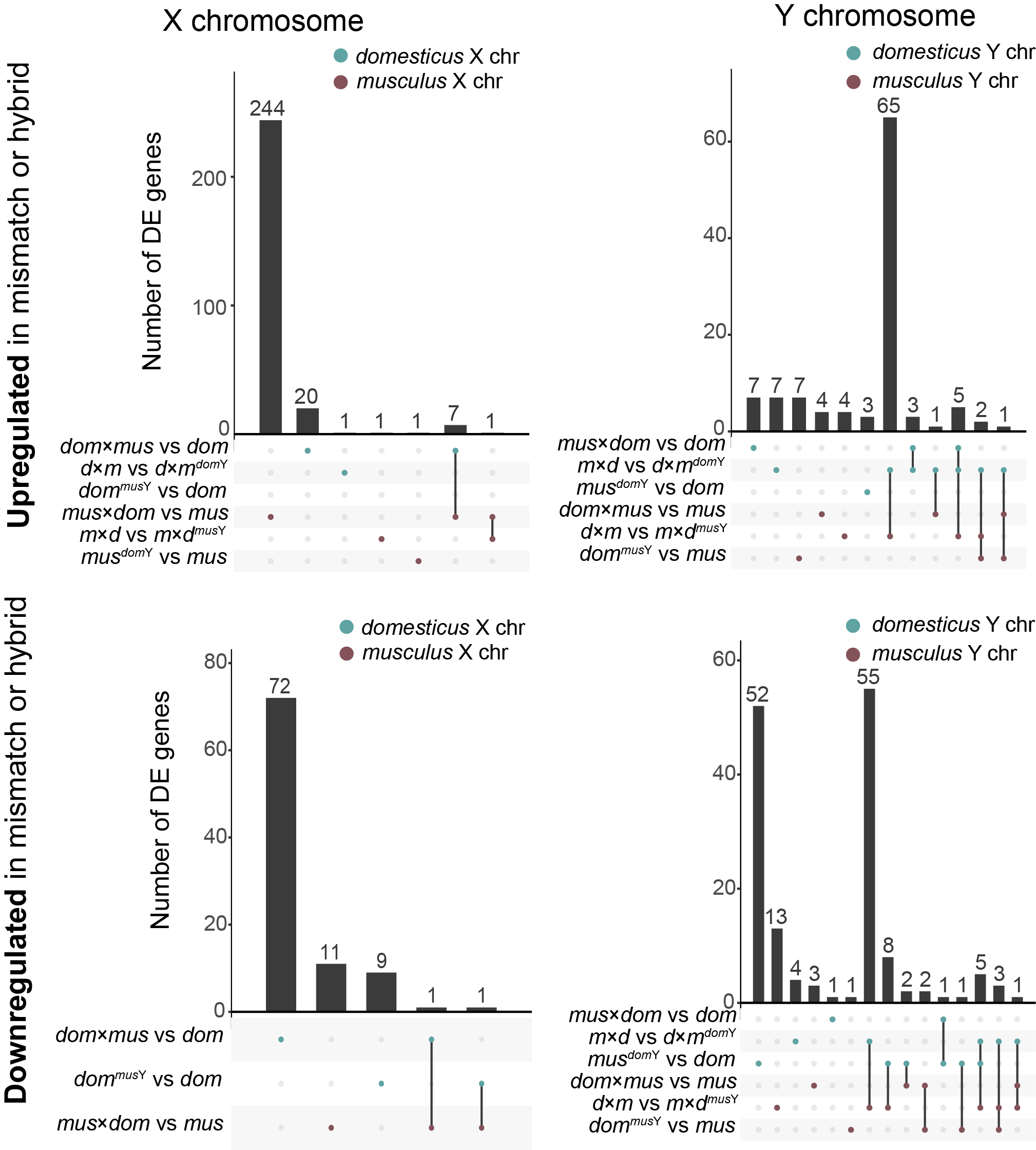

**Figure S7:** Upset plots showing the number of DE genes in each cross type comparison, and genes that are DE across multiple comparisons. (A) DE genes on the X chromosome overexpressed in F1 hybrids or XY mismatch mice relative to controls. (B) DE genes on the Y chromosome overexpressed in F1 hybrids or XY mismatch mice relative to controls. (C) DE genes on the X chromosome underexpressed in F1 hybrids or XY mismatch mice relative to controls. (D) DE genes on the Y chromosome underexpressed in F1 hybrids or XY mismatch mice relative to controls. Bars corresponding to multiple dots connected by lines indicate genes that are DE across multiple comparisons. Bars corresponding to single dots indicate genes that are DE in only one comparison. Blue dots indicate comparisons on the *domesticus* X chromosome (A and C) or *domesticus* Y chromosome (B and D), and red dots indicate comparisons on the *musculus* X chromosome (A and C) or *musculus* Y chromosome (B and D).

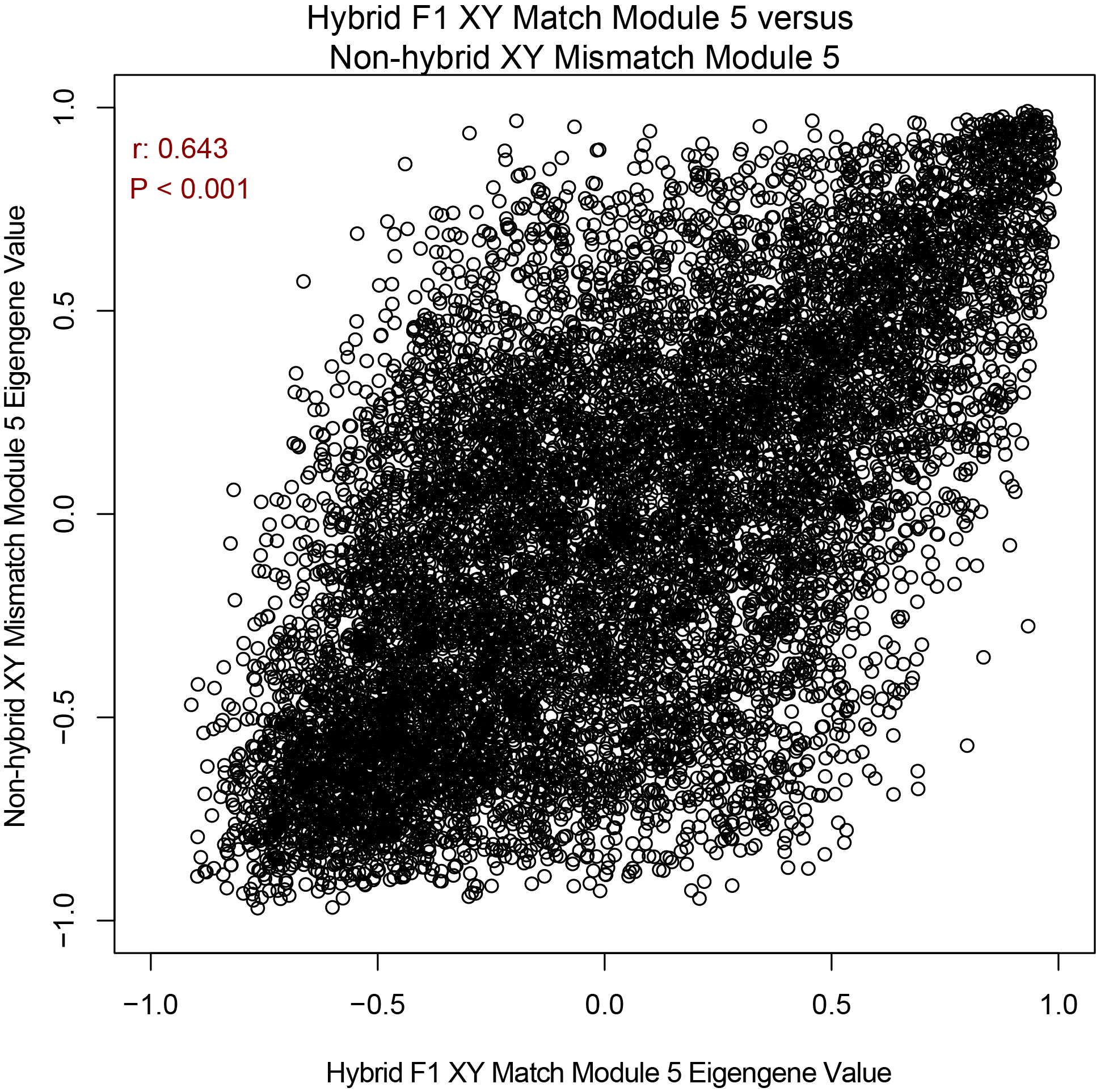

**Figure S8:** Plot showing the correlation between per-gene eigengene value in F1 Hybrid XY Match Module 5 and Non-hybrid XY Mismatch Module 5. Each point represents a gene, with its module membership (module eigengene value) in Hybrid F1 XY Match Module 5 on the x-axis and its module membership in Non-hybrid XY Mismatch Module 5 on the y-axis. Correlation coefficient and p-value are based on a Pearson’s correlation test with FDR correction for multiple tests.

**Supplementary Tables**

**Table S1: RNAseq metadata.** Available as a separate attachment.

LZ = leptotene-zygotene; RS = round spermatid

**Table S2: Male reproductive trait raw phenotype data for each mouse sample.** Available as a separate attachment.

**Table S3: Copy number estimates in wild-derived inbred laboratory strains and Y-introgression strains.** Results are presented using two different methods: a relative coverage approach using Mosdepth (Pedersen and Quinlan 2017) and a k-mer based coverage approach as implemented in AmpliCoNE (Vegesna, et al. 2019). For all genes except *Speer*, paralogs were based on a BLAT search and 97% sequence identity threshold cutoff. For *Speer*, we used a 90% sequence identity threshold cutoff, indicated by (*), because many annotated *Speer* genes had ~90-97% sequence identity with each other based on a BLAT search. Unlike other gene families, *Speer* copy number estimates were very different between the Mosdepth and AmpliCoNE approaches, likely because AmpliCoNE involves a mapping step that requires high sequence identity among paralogs (See Materials and Methods).

|  |  | **Mosdepth** | | | | | | | | **AmpliCoNE** | | | | | | | |
| --- | --- | --- | --- | --- | --- | --- | --- | --- | --- | --- | --- | --- | --- | --- | --- | --- | --- |
| **Cross type** | **Strain** | ***Slx*** | ***Slxl1*** | ***Sly*** | ***Sstx*** | ***Ssty1*** | ***Ssty2*** | ***α-takusan*** | ***Speer**** | ***Slx*** | ***Slxl1*** | ***Sly*** | ***Sstx*** | ***Ssty1*** | ***Ssty2*** | ***α-takusan*** | ***Speer**** |
| *musculus* | PWK | 48 | 34 | 192 | 36 | 136 | 123 | 729 | 238 | 50 | 38 | 213 | 33 | 93 | 135 | 570 | 3 |
| *musculus^dom^*^Y^ | PWK.LY | 50 | 38 | 148 | 40 | 200 | 78 | 729 | 240 | 52 | 35 | 119 | 33 | 147 | 64 | 569 | 3 |
| *domesticus^mus^*^Y^ | LEWES.PY | 17 | 23 | 211 | 45 | 139 | 135 | 259 | 119 | 16 | 21 | 220 | 37 | 80 | 152 | 191 | 3 |
| *domesticus* | LEWES | 16 | 22 | 152 | 43 | 201 | 82 | 254 | 127 | 15 | 20 | 134 | 32 | 160 | 68 | 195 | 3 |

**Table S4: 100kb windows with evidence for introgression.** Available as a separate attachment.

Four introgressed windows (highlighted in gray in the table) were putatively introgressed in both reciprocal Y-introgression strains, which is more than expected by chance (hypergeometric test P << 0.001). These included on region on chromosome 12 and three nearby regions on chromosome 13, one of which contains the gene *Nlrp4f*, which is involved in female fertility (Smith, et al. 2019). Of the putative introgressed regions, 29 windows in *domesticus^mus^*^Y^ and 28 windows in *musculus^dom^*^Y^ were not adjacent to any other window which evidence for introgression, so they likely do not represent long tracks of introgression.

**Table S5: Sex ratios produced by Y-introgression male mice with X-Y mismatch.** Each column represents a different cross type involving Y introgression mice. Note that *domesticus^mus^*^Y,♀^ and *musculus^dom^*^Y,♀^ are females produced from reciprocal backcrosses for generating Y introgression males, but do not have an introgressed Y chromosome because they are females. P-values and chi-squared values are based on a Pearson’s chi-squared test for a significant difference from a 50:50 sex ratio. We did not perform a correction for multiple tests because none of the p-values were significant. Power was calculated based on degrees of freedom = 1 and a significance level = 0.05. Effect sizes for power calculations were calculated by dividing the chi-squared value by the sample size and taking the square root.

|  | *domesticus^mus^*^Y,♀^ × *domesticus^mus^*^Y,♂^ | *domesticus*^♀^ × *domesticus^mus^*^Y,♂^ | *musculus^dom^*^Y,♀^ × *musculus^dom^*^Y,♂^ | *musculus*^♀^ × *musculus^dom^*^Y,♂^ |
| --- | --- | --- | --- | --- |
| **# male offspring** | 40 | 19 | 55 | 29 |
| **# female offspring** | 30 | 24 | 49 | 36 |
| **P-value** | 0.28 | 0.54 | 0.62 | 0.46 |
| **Chi-squared** | 1.157 | 0.372 | 0.240 | 0.554 |
| **Power (1 – Type II Error Probability)** | 0.19 | 0.09 | 0.08 | 0.12 |

**Table S6: Differentially expressed genes on the sex chromosomes and their associated WGCNA modules.** Available as a separate attachment.

Genes with module value “NA” were filtered out of WGCNA based on the *goodSamplesGenes()* function, which checks for factors such as missing data or zero variance. Expression data from Hybrid F1 XY Match and Non-hybrid XY Mismatch were run through WGCNA separately, so some DE genes may belong to two different modules if they were DE in both experiments.

**Table S7: Differentially expressed genes on the autosomes, after removing putatively introgressed regions, and their associated WGCNA modules.** Available as a separate attachment.

Genes with module value “NA” were filtered out of WGCNA based on the *goodSamplesGenes()* function, which checks for factors such as missing data or zero variance. Expression data from Hybrid F1 XY Match and Non-hybrid XY Mismatch were run through WGCNA separately, so some DE genes may belong to two different modules if they were DE in both experiments.
